# *ImplantoMetrics* - Multidimensional trophoblast invasion assessment by combining 3D-*in-vitro* modeling and deep learning analysis

**DOI:** 10.1101/2025.04.25.650556

**Authors:** Ayberk Alp Gyunesh, Marlene Rezk-Füreder, Celine Kapper, Gil Mor, Omar Shebl, Peter Oppelt, Patrick Stelzl, Barbara Arbeithuber

**Author notes:** These authors contributed equally (Ayberk Alp Gyunesh, Marlene Rezk-Füreder). These authors contributed equally (Patrick Stelzl, Barbara Arbeithuber).

## Abstract

Infertility affects millions of couples worldwide, and *in vitro* fertilization is a key therapeutic strategy for achieving parenthood. Despite advances, the first IVF attempt fails in ∼60% of patients, highlighting the need for innovative solutions to improve clinical outcomes. Challenges include the limited ability to study embryo implantation, inadequate methods to test therapeutic drugs, and lack of metrics to evaluate implantation images. To address these issues, we developed *ImplantoMetrics*, a Fiji plugin for quantitative assessment of trophoblast invasion in combination with a 3D-*in-vitro* model. *ImplantoMetrics* uses Convolutional Neural Network and XGBoosting to accurately measure multidimensional expansion patterns. It allows quantitative evaluation of therapeutic interventions, and enables a complex study of trophoblast invasion. Compared to manual methods, *ImplantoMetrics* is ∼13-times faster and reduces errors through automation. Beyond implantation research, *ImplantoMetrics* offers a comprehensive tool to study spheroid invasion in different biological contexts, as e.g. demonstrated here for cancer research.

## Introduction

Infertility is a significant global health issue, with an estimated lifetime prevalence of 17.5% (1). Modern assisted reproductive technologies, such as *in vitro* fertilization (IVF), aim to address this issue. However, live birth rates per IVF cycle remain low: For women under 35 years success rates are about 40% (2), underscoring a critical need for further research in this field. In reproductive research the main emphasis has so far been on enhancing the quality of blastocysts and their implantation (3). Yet, a crucial aspect was frequently overlooked: Low endometrial receptivity is estimated to account for about two-thirds of implantation failures, suggesting that the actual cause of these failures often resides within the endometrium (4). Experimental limitations, such as inadequate animal models for human implantation, present significant challenges in IVF research (5). The adoption of 3D *in vitro* models for human implantation processes has emerged as a promising solution for mimicking these in a laboratory setting (6–8). While advancements have been made, there are still substantial limitations in the methodologies used for quantifying blastocyst implantation *in vitro* (*1,9*).

Different parameters from microscopy images, such as the migration radius and the number of cell projections, were previously measured by manually counting and tracing these features (10,11). This approach, however, is very time-consuming and introduces a potential bias from the person who is performing the analysis. Furthermore, it does not capture additional, potentially critical, aspects such as the distribution of migration or invasion, or the importance of individual parameters over time. Alternatively, the blastocyst area is often measured by manually tracing the shape of the attached blastocyst followed by the determination of the area using Fiji software (a distribution of ImageJ) (7). However, again this method does not account for multiple parameters in parallel, limiting the comprehensiveness and automation necessary for reliable analysis. Moreover, currently used technologies lack the capability to combine multiple parameters, which would be important to fully understand the biology of the implantation process (12). This highlights the need for automated quantification methods to better understand and investigate the complex mechanisms of implantation (13). To bridge the existing gap with advanced quantification, analysis techniques, and practical solutions, we developed *ImplantoMetrics*. *ImplantoMetrics* is a Fiji plugin developed in conjunction with a 3D *in vitro* model for trophoblast invasion, which represents the first automated methodology for the objective quantification and evaluation of this implantation process.

By employing advanced image and machine learning algorithms, *ImplantoMetrics* not only facilitates simultaneous analysis and visualization of image data, but also dynamically adjusts the importance of various features over time, ensuring a nuanced approach to data interpretation. Our tool automates the measurement and interpretation of invasion metrics directly from microscopy images, reduces both time and bias, and adjusts the weighting of different parameters. *ImplantoMetrics* enables researchers to assess e.g., the impact of different drugs on the invasion of trophoblast cells, easily allowing for separate analysis of effects on individual parameters and hence facilitating a deeper understanding of the dynamic interactions in 3D *in vitro* implantation models. In addition to addressing implantation, we could show the versatility of *ImplantoMetrics* in cancer research by analyzing lysosomes in HeLa cell spheroids – demonstrating its capability to analyze dynamic cellular processes in different biological contexts.

## Results

### Establishment of a 3D in vitro model to mimic trophoblast invasion during implantation

The first step was to establish an experimental setting mimicking blastocyst invasion: To overcome ethical considerations and simplify our main approach to develop a novel assessment tool, we focussed on previously published trophoblast invasion models (11). In analogy to former experiments, we introduced a 3D *in vitro* model in the laboratory that would allow the study of early trophoblast invasion by using a combination of three cell types, as detailed below.

1. *Endometrial stromal cells* are pivotal for *in vitro* implantation models but due to scarce adequate human samples and passage limitations of primary endometrial stromal cells in culture medium their experimental usage is challenging (14,15). Therefore, we decided to apply an immortalized telomerase-pretreated human endometrial stromal cell line (HESC), that shows no structural or numerical chromosomal aberrations, in the 3D *in vitro* model (16). As these immortalized HESCs also normally respond to decidualization, we considered those to be appropriate for our model.
2. *Endometrial epithelial cells* represent the epithelial barrier of the endometrium which must be penetrated by the trophoblast to enable successful implantation (17). For similar reasons as mentioned above we used a since decades established human endometrial adenocarcinoma cell line (HEC-1-A) (18,19) for our *in vitro* model. HEC-1-A, which were originally derived from endometrial G2 carcinoma of a hysterectomy sample from a 71-year-old female, have typical epithelial properties and express key implantation-related markers (18,19).
3. *First trimester trophoblast cells* that are capable of mimicking the outer layer of the blastocyst – referred as trophoectoderm – are essential for an adequate experimental implantation *in vitro* setting (20). Therefore, we integrated a telomerase immortalized trophoblast cell line (Sw.71), which was originally isolated from a 7-week old normal placenta, into our model (21). This cell line has similar characteristics to primary first trimester trophoblast cells (21) and can be cultured to form three-dimensional spheroid structures – blastocyst-like spheroids (BLS) – which mimic the trophectoderm (11).

*<u>Experimental Design:</u>* The cell preparation and transfer process extended over a period of four days, followed by imaging from day four to day ten. HESCs seeded on day one represent the stromal cell layer of the endometrium. On day two, HESCs were covered with Matrigel, which was used to simulate the human endometrial extracellular matrix. After solidification, HEC-1-A cells were added on top, representing the epithelial layer of the endometrium (***Fig 1A***). In parallel, BLSs were prepared from green fluorescent protein (GFP) labeled Sw.71 trophoblast cells in ultra-low attachment plates, ensuring their spherical development over two days (11). On day four, BLS was transferred to the cell layer structure that mimics the endometrium, and with or without drug addition, images of each well were captured at 8-hour intervals using a cell imaging multimode reader (Cytation 7) or at 12-hour intervals using an inverted fluorescence microscope (Olympus IX73). Image acquisition continued for up to six days and the obtained images were used to quantify the invasion process using the *ImplantoMetrics* plugin for Fiji (***Fig 1A***), a tool we developed that enables automated, robust, and standardized analysis of invasion metrics. The high numbers of images (and hence individually monitored BLS invasion events) required for tool development necessitated us to keep our endometrial model simple. However, *ImplantoMetrics* can also be applied in conjunction with more complex settings (e.g., models that include decidualized cells, more complex layered endometrial models, or endometrial organoid models that allow for trophoblast invasion imaging (e.g., 7,22)).

**Figure 1.**
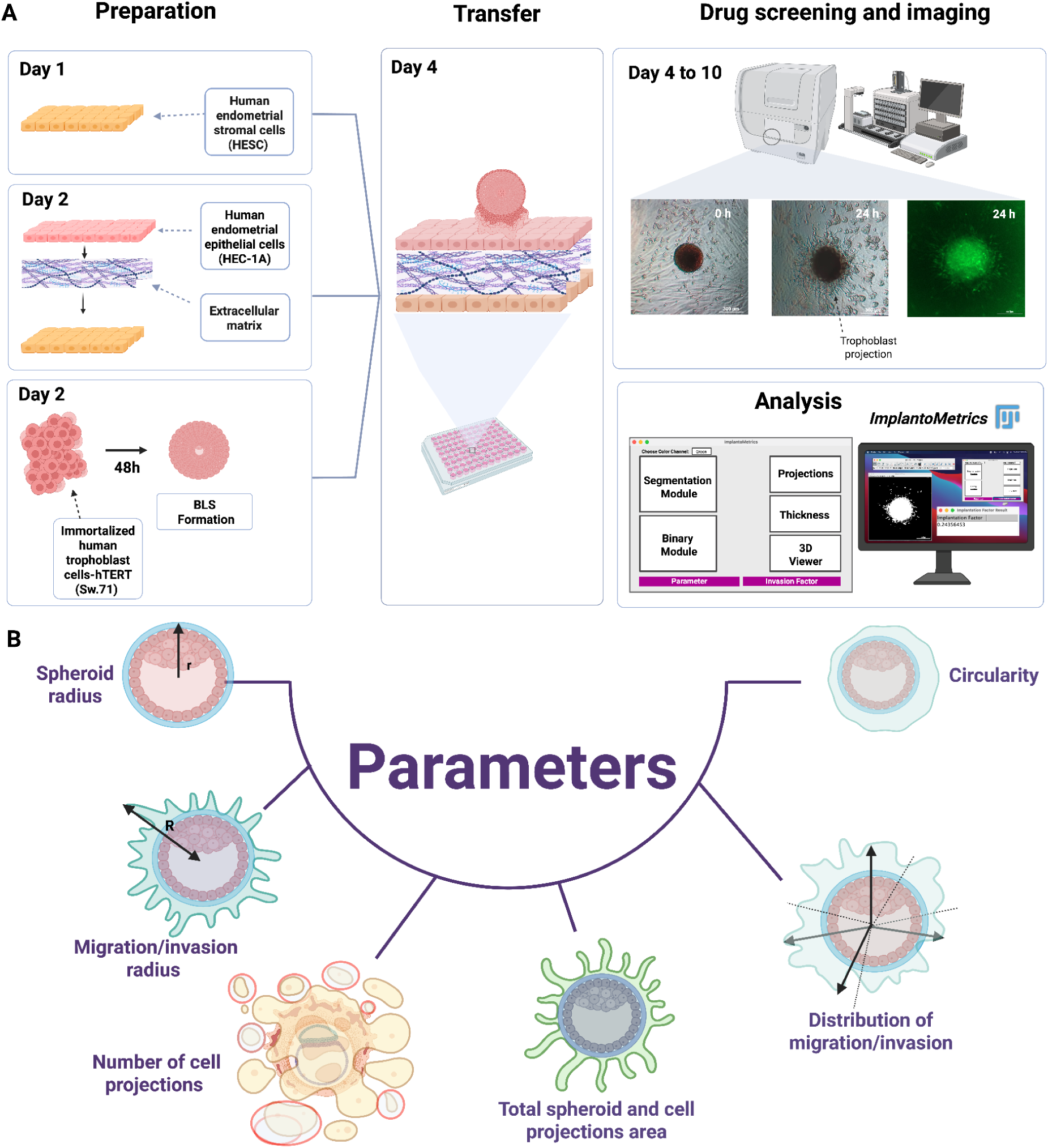
3D *in vitro* model to study BLS invasion, assessed by *ImplantoMetrics*. **A)** On day one, HESC cells were seeded into a 96-well plate. On day two, a Matrigel layer was applied and HEC-1-A cells were added on top. In parallel, BLSs were grown from Sw.71 cells in a separate ultra-low attachment 96-well plate. After two days, the BLS developed a spherical structure. On day 4, the BLS was transferred to the cell layer model, placing the BLS in the center of the wells. BLS invasion was monitored for up to six days. During that period, the effect of different drugs on BLS invasion can be tested. **B)** *ImplantoMetrics* was used to analyze the invasion process and it gives six parameters that are important in the assessment of invasion success: spheroid radius, total spheroid and cell projections area, migration/invasion radius, number of cell projections, distribution of migration/invasion, and circularity. Each parameter has a different level of significance in implantation assessment and influences the process differently over time.

### *ImplantoMetrics* - A tool to assess spheroid invasion *in vitro*

*ImplantoMetrics* is a deep image analysis tool based on Convolutional Neural Networks (CNNs), that provides a new, automated method to identify and quantify cell invasion metrics in *in vitro* implantation experiments (***Fig 1A***). The input data are microscopic images of fluorescently labeled spheroid structures (such as BLSs), from which six parameters are extracted: spheroid radius, migration/invasion radius, number of cell projections, total area of the spheroid and cell projections, distribution of migration/invasion, and circularity (***Fig 1B***). These parameters were chosen based on their critical roles in assessing cell behavior and invasion dynamics in spheroid models and by providing a comprehensive overview of the structural and spatial characteristics of spheroids (23–26). For instance, various implantation-associated gene expression changes in the trophectoderm were shown to be related to morphological changes (27,28), and are considered as key indicators of trophoblast differentiation and implantation potential (28). Additionally, lamellipodial protrusions are crucial for embryo–endometrium interactions (28). Moreover, parameters such as spheroid size, shape, and circularity reflect shifts from proliferative to invasive states, highlighting differences between polar and mural trophoblast cells critical for establishing maternal contact zones (28).

By focusing on these morphological indicators—including migration/invasion radius, number of trophoblast projections, circularity, and migration/invasion distribution—*ImplantoMetrics* can quantitatively assess various processes essential for trophoblast attachment and successful implantation. *ImplantoMetrics* further offers integrated visualization methods such as cell projections, thickness, and a 3D viewer for detailed visualization and analysis of the invasion process. An ‘*Invasion Facto*r’ (ranging from zero to one) is calculated to quantitatively assess the probability of successful invasion of trophoblast cells based on the predicted parameters and extracted image features.

#### CNN training to generate a model for automated parameter determination

CNNs are used to enable automated and accurate determination of the six parameters (spheroid radius, total spheroid and cell projections area, migration/invasion radius, number of cell projections, distribution of migration/invasion and circularity) from microscopic images of fluorescently labeled spheroids. Within the CNN architecture, *ImplantoMetrics* uses the Xception model as the base model, which is known for its efficiency and accuracy in handling complex image data (29). Xception has shown superior performance in various image classification and regression tasks compared to models such as VGG16, ResNet, and Inception during transfer learning (based on Hyperopt analysis, see ***Methods*** for details; ***S1 Fig, S1 Table***). This is primarily because Xception employs depthwise separable convolutions, which reduce the computational cost while maintaining high accuracy (30), which makes it particularly suitable for processing the high-dimensional and complex image data used in our study. To increase data variance and minimize overfitting, data augmentation techniques such as image rotation, shearing, and mirroring were used (***S2 Table***) (31). Furthermore, the integration of batch normalization contributes to stabilizing and accelerating the learning process by minimizing internal covariate shifts and thus increasing training efficiency (32), and other hyperparameters (based on Hyperopt analysis, see ***Methods*** for details), including L1-L2 regularization to avoid overfitting and dynamic adjustment of the learning rate using ‘ReduceLROnPlateau’ and ‘EarlyStopping’ to terminate training in the absence of improvements in validation losses, play important roles (***S2 Table***) (33,34). Min-max normalization was used to standardize the measured data and outliers were removed using the interquartile range (IQR) method.

A total of 1,872 images of BLS invasion (divided into 80% training, 10% test, and 10% validation datasets, see ***Methods*** for details) were used to develop our model. In the pre-processing phase, Segmentation module and Binary module were employed to segment the input images and create binary masks ***(Fig 2AB***, *see **Methods** for details)*. The binary images were expanded to three channels to ensure compatibility with the Xception model, which was loaded without the top layers, but with a customized input layer that can process single-channel images. This adaptation was done by a lambda layer that extends the single-channel images to three channels **(*Fig 3A*)**. The model’s output features were then processed through additional layers: First GlobalAveragePooling2D reduced the dimensionality of the feature maps. This was followed by Dense layers with 2,048 and 256 neurons, using ReLU activation, L1-L2 regularization, and BatchNormalization to stabilize the learning process. The final output layer consisted of six neurons with a linear activation function, representing the six parameters to be predicted (spheroid radius, total spheroid and cell projections area, migration/invasion radius, number of cell projections, distribution of migration/invasion, and circularity). The exact measurement methods of the six parameters and formulas are detailed in ***S2-6 Figs***.

**Figure 2.**
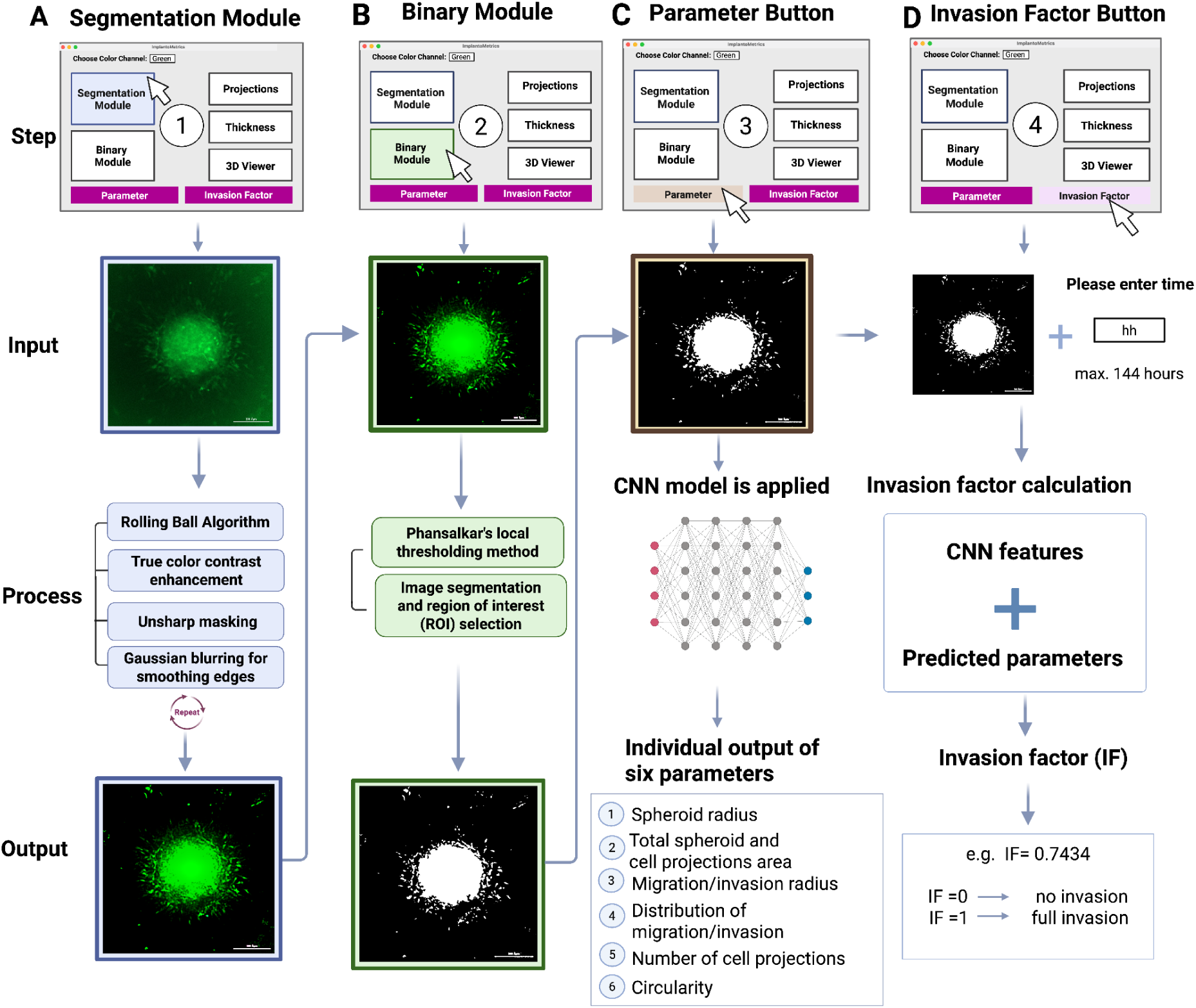
Image preprocessing and algorithm integration. **A)** *ImplantoMetrics* analysis starts with the selection of the ‘Segmentation Module’ to initiate the segmentation of the image content. After successfully completing the segmentation, the interface will indicate that the process is complete. In cases of unsuccessful segmentation, a feedback message for troubleshooting is displayed, indicating required image properties for successful segmentation. **B)** The Binary Module is selected and the image is then converted into a binary format, which facilitates further analysis. **C)** The ‘Parameter’ button is used to upload the *ImplantationMetrics* model, which measures six parameters: spheroid radius, total spheroid and cell projections area, the migration/invasion radius, the distribution of migration/invasion, the number of cell projections and circularity. **D)** Finally, the user clicks the ‘*Invasion Factor*’ button and enters the duration of the invasion process in hours (’hh’ format) to calculate the invasion factor, which quantifies the probability of successful trophoblast invasion based on the data entered and the calculation of the model.

**Figure 3.**
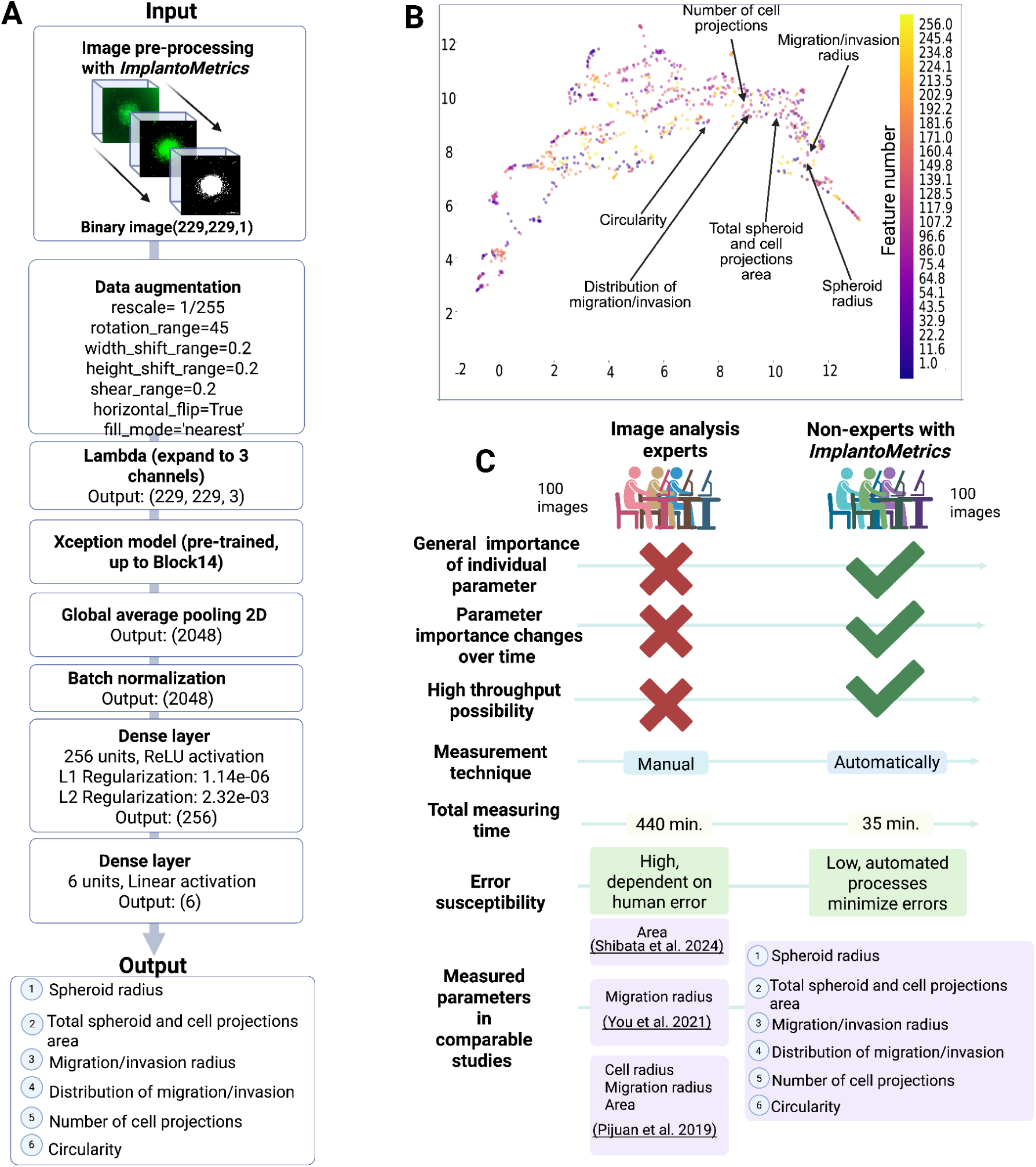
CNN Model structure, training process, and UMAP (Uniform manifold approximation and projection) analysis. **A)** Machine learning framework. Initially, an image processing plugin segments images, which are then used as input for a CNN. The architecture includes data augmentation, dimensional expansion, a pre-trained Xception model up to Block14, global average pooling, batch normalization, and dense layers. The final output layer uses ReLU and linear activations to predict various metrics such as spheroid radius, total spheroid and cell projections area, migration/invasion radius, distribution of migration/invasion, number of cell projections and circularity. **B)** UMAP projection is used to visualize the relationships and patterns between different invasion parameters (n=6) and extracted features (n=256). The axes, UMAP 1 and UMAP 2, represent the reduced dimensions derived from a variety of measured features. Each point on the map corresponds to a specific parameter or feature. The color scale from yellow to purple on the right represents the feature number, with each color assigned to a different feature number. **c)** Comparison of manual image analysis by experts with non-experts using *ImplantoMetrics*

To visualize the relationships between the invasion parameters and all features before the final output layer, we used uniform manifold approximation and projection (UMAP). The UMAP projection revealed clusters and separations among the parameters, which provides insights into the complex interactions relevant to trophoblast invasion **(*Fig 3B*)**. After the training phase, the model was converted with the tf2onnx library to the Open Neural Network Exchange (ONNX) format. The use of ONNX Runtime facilitates the implementation and scaling of CNNs in medical image processing by enabling consistent execution of models across different platforms (35).

#### Evaluation of parameter/feature-importance using the XGBoost algorithm

We next wanted to examine the temporal dynamics and significance of individual features in the invasion process by employing the eXtreme Gradient Boosting (XGBoost) algorithm, (which emerged as the most effective algorithm, ***S3 Table***). We utilized our dataset of 1,872 BLS invasion images to develop two models: Model A, which used the measured six parameters from the images as input, and Model B, which used CNN-extracted features as input (***Fig 4A-D***). For both models, the output was the predicted “time” to calculate the temporal importance of the parameters/features using SHapley Additive exPlanations (SHAP) values. The optimized hyperparameters for XGBoost models A and B were determined using randomized search and the models were trained with these best-found hyperparameters. The combined model, including the validated metrics reflecting the accuracy of the models, is listed in ***S3 Table***. The significance of each parameter/feature for the prediction was quantified as SHAP values, which reflect the influence of individual parameters/features within the model’s context for Model A and Model B **(*Fig 4E-H*)**: Each SHAP value quantifies how the inclusion or exclusion of a specific parameter affects the model’s prediction, drawing upon cooperative game theory to equitably distribute the influence of each feature.

**Figure 4.**
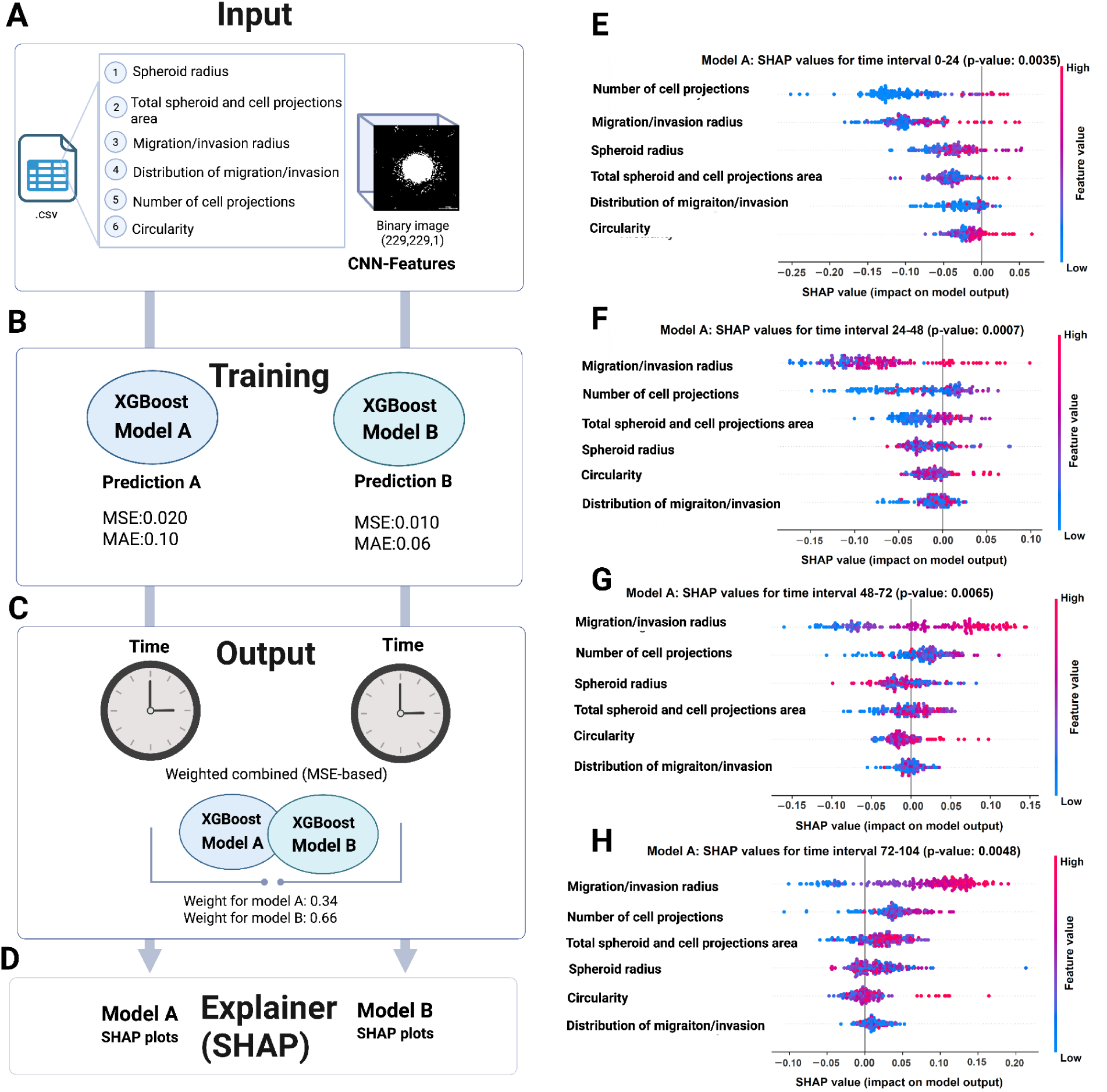
Integration of XGBoost and SHAP to quantify the probability of invasion. **A**) Pre-processing architecture of our model, showing the input phase where the six measured parameters and significant features from a CNN are fed into XGBoost Model A and Model B, respectively. **B**) Training and validation phase for XGBoost Model A and Model B, the processes of Randomized Search optimization and the evaluation loop involving mean squared errors (MSE) and Mean Absolute Error (MAE) metrics. **C**) Ensemble formation phase, in which the predictions of XGBoost Model A and Model B are combined to improve the predictive performance of the overall model. The output is the predicted ‘’Time’’. The combination of the models is based on the weighted MSE, with Model A receiving a weight of 0.34 and Model B a weight of 0.66. **D)** SHAP value analysis results from our XGBoost model, which predicts the timing of BLS invasion across four distinct time intervals. Intervals are: **E)** 0-24 hours, **F)** 24-48 hours, **G)** 48-72 hours, and **H)** 72-104 hours (after 104 hours, no significant changes were measured). Each subsection of the graph depicts the average SHAP values of the parameters within a given interval, with statistical significance determined using Welch’s t-tests (p-value ≤ 0.05). Horizontal axes represent the SHAP values, indicating the impact of each parameter on the model’s prediction for the timing of invasion. Positive SHAP values suggest that the parameter contributes to a later invasion time, whereas negative values indicate an association with an earlier invasion time. The vertical axis lists the parameters under investigation. Red and blue dots illustrate the parameter levels: for instance, an increase in the migration/invasion radius (red) correlates with a positive influence, while a shift towards blue signifies a more negative impact.

Building on this, we computed the aggregate of the SHAP values across all observations to grasp the overall strength of each parameter’s effect. This approach ensures that our analysis is not skewed by outliers or moments that are not representative of the general trend. Consequently, it allows us to pinpoint which parameters/features consistently play a substantial role in the model’s predictions. By comparing the SHAP values at 8-hour intervals we can determine if and when the significance of each parameter/feature changes over time. We visualized the results for the time intervals where significant differences in SHAP values were found (e.g., 0-24 hours, 24-48 hours, 48-72 hours, and 72-104 hours, as shown in ***Fig 4E-H, S7 Fig***; The number of cell projections e.g. had the greatest impact on the predicted outcome in interval 0-24h).

#### Quantification of the trophoblast invasion probability using the invasion factor

Using the analysis of SHAP values and the determined six parameters, *ImplantoMetrics* calculates the ‘*Invasion factor*’, implemented to quantitatively assess the probability of successful trophoblast invasion in a given experiment. By employing the sigmoid function σ(𝐼𝑛𝑣𝑎𝑠𝑖𝑜𝑛 𝐹𝑎𝑐𝑡𝑜𝑟) any real number is mapped into a bounded interval from zero to one, which is suited to the nature of probability predictions in our regression problem. While the sigmoid function is known for its application in binary classification, its utility extends to regression tasks where the output is probabilistic (36). In our context, the sigmoid function ensures that the ‘Invasion factor’ remains in the interpretable range of zero (indicating very low probability of trophoblast invasion) to one (indicating very high probability of trophoblast invasion), regardless of the magnitude of the input.

The ‘*Invasion factor*’ is calculated as:

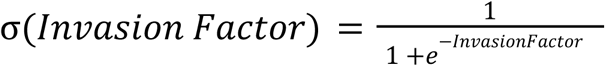

Where the ‘*Invasion facto*r’ is defined by :

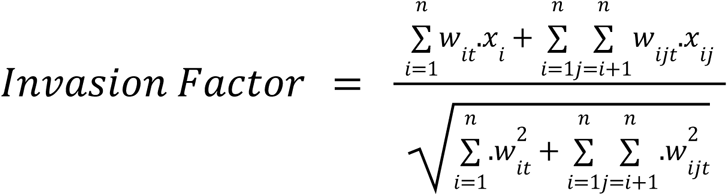

Here, 𝑤_𝑖𝑡_ denotes the SHAP values for the 𝑖-th feature at time 𝑡, capturing both its positive and negative influences on the model’s output ‘’Time’’. 𝑤_𝑖𝑗𝑡_ signifies the SHAP value for the interaction effect between the 𝑖-th and 𝑗-th features at time 𝑡. 𝑥_𝑖_ is normalized value for the 𝑖-th feature, ensuring that feature values are on the comparable scale. 𝑥_𝑖𝑗_ represents the interaction term, obtained by multiplying the normalized values of feature 𝑖 and 𝑗.

Each characteristic is multiplied by its corresponding SHAP value, which quantifies its importance in the predictive model. These products are then aggregated to form a composite score. This score is normalized by dividing it by the sum of the average SHAP values of all characteristics, creating a weighted average. It also maintains the sensitivity to the magnitude of the SHAP values and the directionality of each feature’s impact, whether positive or negative. The numerator of the formula sums up all individual and interactive contributions from the features, while the denominator normalizes these contributions. The resulting ‘’*Invasion factor*’’ serves as a comprehensive metric, encapsulating the combined influence of all characteristics in a balanced way.

### Usage of the *ImplantoMetrics* plugin

The Fiji plugin *ImplantoMetrics* can be downloaded from GitHub under the following link: https://github.com/creativebrain1729/ImplantoMetrics/releases/tag/v.1.0.0. A detailed description on how *ImplantoMetrics* can be downloaded and installed is shown in ***S8 Fig*** and also available in the respective GitHub repositories along with sample images. When using the *ImplantoMetrics* plugin, first the color channel for analysis needs to be specified (the default color channel is green). Next, the ‘Segmentation Module’ needs to be selected for fluorescence image segmentation (***Fig 2A*)**. The module then performs a series of image processing algorithms, including the rolling ball algorithm for background correction, true color contrast enhancement to improve the contrast, application of unsharp masking to sharpen the image, and Gaussian blurring to smooth the edges (*see **Methods** for more details*). These steps are automatically repeated four times to optimize the quality of the image segmentation. When segmentation is finished, the image undergoes an automated quality assessment which determines whether the image meets the required quality standards for successful segmentation (***S9-11 Figs***). This assessment is performed using a combination of Receiver Operating Characteristic (ROC) analysis and histogram-based thresholding, evaluating key quality metrics such as contrast, entropy, homogeneity, and correlation. If the image meets the threshold criteria, the confirmation message “*Segmentation was successful*” appears. If the image does not meet the quality criteria, the system provides a feedback message for troubleshooting, indicating which quality metrics are suboptimal and suggesting possible improvements (***S9 Fig***).

Once a validated, segmented image is available, the ‘Binary Module’ can be selected, which uses the fluorescent channel of the provided image as input (***Fig 2B***). By applying Phansalkar’s local thresholding method, the image is binarized (converted into a two-color format). This step facilitates the identification and delineation of regions of interest (ROI) in the image and is crucial for the efficiency of subsequent model processing steps (***Fig 2B*** *see **Methods** for details*). The binarized image clearly distinguishes the structures of interest from background and is thus used for subsequent analysis steps (***Fig 2B*)**. Next by clicking the ‘Parameter’ button, the ‘*ImplantationMetrics* model’, which was trained as described before in ‘CNN training to generate a model for automated parameter determination’ is uploaded. This model is used to predict six parameters that can vary depending on invasion success (***Fig 1A*, *Fig 2C***). The result of this analysis is then output individually for each of the six parameters, providing a detailed profile of the investigated structure for further scientific evaluation or data presentation. Finally, by selecting the ‘*Invasion factor*’ button and entering the acquisition time of the image, the ‘*Invasion factor*’ is calculated based on the predicted six parameters and extracted CNN features from the image, representing the probability of successful invasion based on the specified image and time (***Fig 2D***).

### ImplantoMetrics visualizations

The *ImplantoMetrics* plugin interface also offers three integrated visualization methods: Projections, Thickness, and 3D Viewer. The ‘Projections’ button allows the visualization of the distribution and appearance of the spheroid and its extensions. The spheroid is highlighted in red and projections in green (***Fig 5A***). The Thickness button allows a 3D visualization of the cell thickness/diameter, as well as an intensity distribution of the image, which can be used to identify areas of potential interest (***Fig 5B***). If Z-stack images of different time intervals are available, the 3D Viewer can be used to generate a 3D visualization, providing more insight into the invasion process (***Fig 5C***).

**Figure 5.**
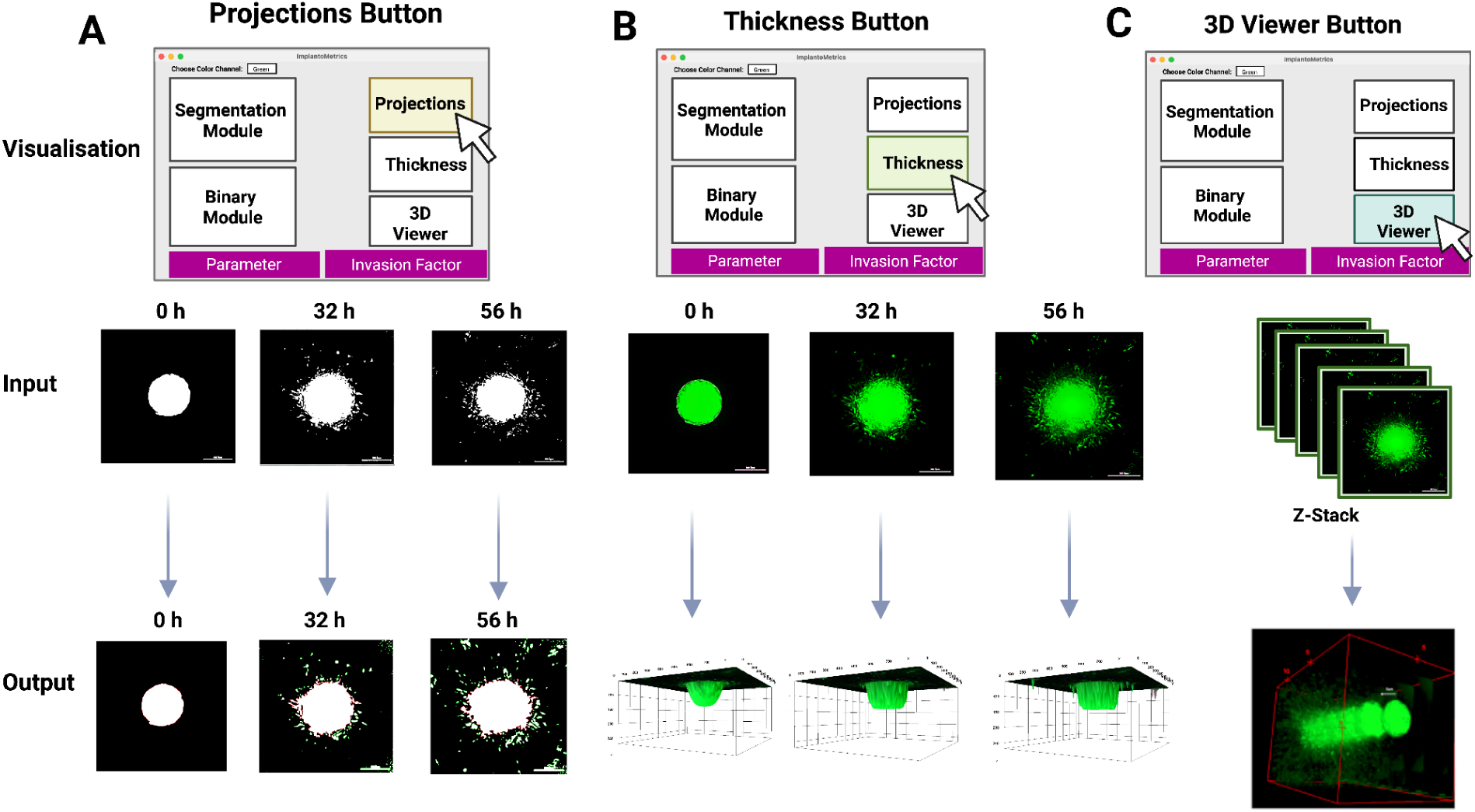
*ImplantoMetrics* analysis visualization. **A)** The ‘Projections’ feature allows visualization of projections and BLS, with red highlighting BLS for identification and green indicating projection locations, aiding in assessing cell distribution. **B)** Thickness Button offers a 3D Surface Plot for cell thickness for intensity distribution, useful for evaluating implant quality and pinpointing potential issues. **C)** When Z-stack images from various time frames are available, a 3D representation can be generated, enhancing the visualization’s depth.

### Comparison of *ImplantoMetrics* to manual invasion analysis

We demonstrated the efficiency and accuracy of *ImplantoMetrics* compared to manual measurement by conducting a comparative analysis (***Fig 3C***): Experts (n=3) manually analyzed 100 images, taking on average 440 minutes (SD = 8.16 minutes) and yielding results prone to human error. In contrast, *ImplantoMetrics* enabled non-experts (n=3) to analyze the same number of images 13-times faster (on average 35 minutes, SD = 6.24 minutes). Additionally, *ImplantoMetrics* provides a consistent and reproducible analysis by minimizing human error and bias through automation, allowing for a high-throughput and dynamic evaluation of multiple parameters (***Fig 3C***). The separate assessment of parameters and the calculation of the ‘*Invasion facto*r’ provide valuable insights into the dynamics of the implantation process which is hard to address in manual image analyses.

#### *ImplantoMetrics* reveals successful inhibition of BLS invasion upon indomethacin treatment

To validate the performance of the ‘*Invasion factor*’ in predicting trophoblast migration/invasion outcomes, we used the before described 3D *in vitro* endometrial model (***Fig 1A***) with or without the addition of indomethacin, a drug that inhibits trophoblast invasion: Indomethacin is known to impact the production of for implantation critical biomolecules such as cytokines, matrix metalloproteinases (MMPs), tissue inhibitors of metalloproteinases (TIMPs), and prostaglandin E2 (PGE2) (37). In addition to our combined model (consisting of HESCs, Matrigel, and HEC-1-A), we also tested the impact of indomethacin (applied at three different concentrations) on BLS invasion in HESCs with Matrigel (hence lacking of any potential cancer associated bias from the adenocarcinoma-derived cell line HEC-1-A) or HEC-1-A cells with Matrigel. Images were taken at 8-hour intervals using a cell imaging multimode reader (Cytation 7) and analyzed with *ImplantoMetrics* to calculate ‘*Invasion factor*’ ***(Fig 6***). We could show that without indomethacin, the combination of HESC and HEC-1-A cells as endometrial model leads to earlier and more efficient invasion of trophoblast cells compared to HESC or HEC-1-A cells alone (reaching an *Invasion factor* of ∼0.85 after 80h for HESC+HEC-1-A; ***Fig 6A***), whereby HESC cells (which reach an *Invasion factor* ∼0.85 only after 96h; ***Fig 6B***) show more efficient invasion compared to HEC-1-A cells (which only reach a maximum *Invasion factor* ∼0.80; ***Fig 6C***). In all endometrial models, invasion of trophoblast cells could be efficiently inhibited with indomethacin, resulting in rather constant *Invasion factors* around 0.6. For HEC-1-A cells, however, a significant inhibition of invasion was only observed at the highest concentration of indomethacin (0.6 µg/ml; ***Fig 6C***). To also test the performance of *ImplantoMetrics* using a different image acquisition system, we repeated the experiment taking images at 12-hour intervals (up to 36h) with an inverted fluorescence microscope (Olympus IX73). Again, a significant inhibition of trophoblast cell invasion could be measured (***S12 Fig***).

**Figure 6.**
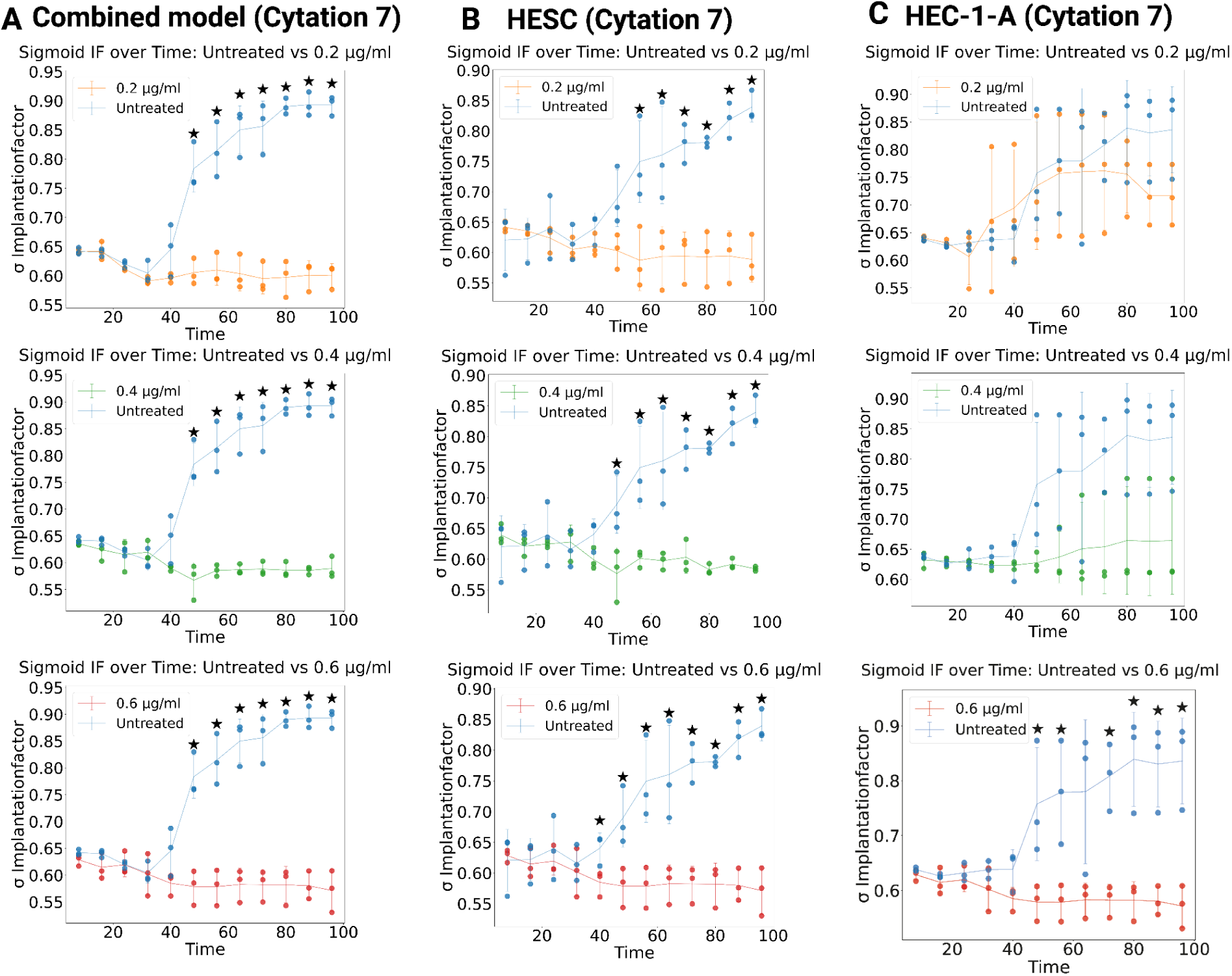
Inhibition of BLS invasion by indomethacin treatment. The mean *Invasion factor* (σIF) (n=3, distinct samples) at three different indomethacin concentrations (0.2, 0.4, and 0.6 μg/ml) is shown for three different *in vitro* models, measured every 8 hours with the Cytation 7: **A)** combined endometrial model (HESC with Matrigel and HEC-1-A), **B)** HESC with Matrigel, and **C)** HEC-1-A with Matrigel. Error bars represent the standard deviation (± SD). Asterisks indicate significant differences (Welch’s t-test, one-sided) between untreated controls and indomethacin treated cells.

### Improved trophoblast invasion upon TNA-α treatment quantified with the *Invasion factor*

We next assessed the ability of the *Invasion factor* to quantify enhanced trophoblast invasion by tumor necrosis factor-alpha (TNF-α) treatment. TNF-α is a well-characterized modulator of trophoblast invasion, regulating key biomolecules such as MMPs, adhesion molecules, and inflammatory mediators (38–40). Beyond its role in ECM remodeling, TNF-α modulates the endometrial stromal secretome, establishing a microenvironment that enhances trophoblast motility (17). Additionally, TNF-α has been shown to upregulate ICAM-1 expression in endometrial epithelial cells, promoting adhesion and implantation processes (40). We applied three different TNF-α concentrations (10 ng/ml, 25 ng/ml, and 50 ng/ml) to our combined model (HESCs, Matrigel, and HEC-1-A). Images were taken at 8-hour intervals and analyzed with *ImplantoMetrics* to calculate the *Invasion factor* (***S13 Fig***). While 10 ng/ml TNF-α did not show a significant difference (***S13A Fig***), 25 ng/ml resulted in a significant increase in trophoblast invasion from 75 hours onward (***S13B Fig***). The strongest effect was observed with 50 ng/ml TNF-α, where trophoblast invasion significantly increased from 56 hours onward (***S13C Fig***). Our results show that TNF-α exerts a concentration dependent effect, with significant invasion enhancement observed at 25 and 50 ng/ml. These findings are consistent with previous reports demonstrating that TNF-α plays a pivotal role in trophoblast invasion by regulating ECM remodeling and cell adhesion (17,39), while also modulating endometrial secretory factors that contribute to implantation success (40).

### Application of *ImplantoMetrics* in embryo implantation-independent 3D *in vitro* experiments

Despite using a model that was trained for BLS invasion, to demonstrate the broader usability of *ImplantoMetrics*, we analyzed lysosomal dynamics (using previously published confocal laser scanning microscopy images) in HeLa cell tumor spheroids during apoptosis, stained with the fluorescent probe L-lyso which can be used to track lysosomes in live cells (41). Analogous to BLS analysis, we quantified the following parameters (not all of the six parameters relevant for implantation play an important role in this HeLa spheroid system): lysosomal number (equivalent to cell projections), circularity of the lysosomal distribution within the spheroid, and total lysosomal area *(**S14 Fig**)*. We could show a positive correlation between the total lysosomal area and the number of lysosomes, indicating that a larger overall lysosomal volume is accompanied by an increase in lysosome number and not a larger size of individual lysosomes. Additionally, a positive correlation was observed between the number of lysosomes and their circular distribution within the spheroid. Conversely, larger spheroids showed a slight tendency to have a less circular distribution of lysosomes. Interestingly, in the one-photon fluorescence images of the 3D HeLa tumor spheroid incubated with L-lyso (60 µM for 1h), circularity fluctuated across the spheroid, with an initial decrease (accompanied by a decline in the number of lysosomes) followed by a substantial increase (accompanied by an increase in the number of lysosomes), indicating morphological changes during apoptosis.

## Discussion

*In vitro* models are crucial for the investigation of the embryo implantation processes and have been heavily utilized in the field (reviewed in 8). Yet, the critical aspect of their quantification remains mainly overlooked: Often, quantification relies on either analyzing single parameters – like basic area measurements with Fiji (7) – or using subjective methods that are prone to be biased, such as manual counting (17,42), neither of which capable of capturing the complexity of the implantation processes. To address this issue, we developed *ImplantoMetrics*, a tool that automatically quantifies multiple parameters relevant to the invasion process, assisted by both machine learning and deep learning for the assessment and quantification. We used CNNs as they are particularly powerful in medical imaging: They are able to recognize subtle and complex patterns in data which are often missed in manual analyses (43). The XGBoost algorithm, which we used in combination with CNNs, was previously shown to be particularly effective in modeling complex nonlinear patterns and is therefore instrumental in identifying important variables in predictive models (44). We therefore used the XGBoost algorithm to analyze the temporal dynamics of the trophoblast cell invasion process in detail, as well as to determine the importance of individual parameters or features. This ensures that the investigation is focused on the influential parameters, which simplifies and refines the biological interpretation of the data and furthermore enables us to not only identify essential parameters, but also to analyze their critical periods of influence. With SHAP swarm plots we show changes in the significance of individual parameters and features over different time intervals of the captured trophoblast invasion process. SHAP analysis is widely applied in clinical settings such as embryo selection and viability prediction, and highlights the importance of identifying parameters that drive biological outcomes (45). For instance, studies using SHAP models to predict clinical pregnancy outcomes emphasize that integrating explainable AI methods can effectively reveal parameter importance, providing insight into underlying biological processes (45,46). Using this feature of *ImplantoMetrics*, we could show for example that the number of cell projections had a greater impact on invasion success within the first 24 hours (reflecting early cytoskeletal activity), while in later intervals (marking the expansion phase of invasion), the migration/invasion radius becomes more significant **(*Fig. 4E-H*).** This finding underscores the high dynamics of, and the temporal changes in morphological parameters during the implantation process, which have been reported previously (28,47,48).

Furthermore, with the implementation of the *Invasion factor* as a quantitative measure for trophoblast invasion success in cell culture based *in vitro* experiments, we enable an easier interpretation and comparability of individual experiments, especially when e.g. the effect of different drugs is tested. As there are no comparable quantitative methods to date, this approach fills an important gap in research. In our 3D *in vitro* model we can dynamically investigate cellular interactions between the endometrial and trophoblast cell lines. Previous research has demonstrated the effectiveness of BLS in mimicking the trophectoderm, the outer layer of the blastocyst (which is critical for implantation), overcoming the ethically challenging need of human embryos (49). The selection of the Sw.71, HEC-1-A, and HESC cell lines for our study is based on their essential role in cell culture based implantation research, mimicking important features of the blastocyst and the endometrium (50–52). The efficacy of each of these cell lines in relation to their specific role in the implantation process has already been demonstrated: Sw.71 cells, known for their HLA-C expression, are particularly relevant for understanding immune tolerance in the context of implantation (50). Immortalized HESCs, derived from endometria of women within the reproductive age (obtained from hysterectomies for benign conditions), have proved to be a valuable resource for basic implantation studies (14–16). HEC-1-A cells are pivotal for understanding the interactions between the embryo and the epithelial layer of the endometrium (52). By comparing the BLS migration/invasion process in an endometrial system consisting of either stromal cells in combination with extracellular matrix and epithelial cells, or either of the two cell types alone in combination with extracellular matrix, we could quantify the dynamics of trophoblast migration/invasion in dependence of cell-type specific signaling. We demonstrated that both cell lines influence BLS invasion, which is in agreement with previous reports (53,54). Treatment with indomethacin inhibited the invasion process in a dose-dependent manner, confirming the previously reported role of cytokines and matrix metalloproteinases, among others (55), while TNF-α enhanced trophoblast invasion reflected by an increase in the *Invasion factor*.

In general, our method opens new avenues for understanding human implantation processes, potentially leading to the development of novel therapeutic interventions in association with implantation failure, and enhancing the overall understanding of reproductive medicine.

However, it is also crucial to acknowledge the limitations inherent in our study due to its *in vitro* nature. The challenges of studying implantation in living organisms, due to ethical and technical issues, require the use of *in vitro* models. While cell lines represent an important tool in implantation research by overcoming several limitations associated with the use of primary tissues (such as limited sample material), they also harbor various disadvantages. For instance, HEC-1-A cells were derived from endometrial adenocarcinoma and therefore add potential cancer-related bias to the system. Furthermore, because we focused on some of the most prevalent cell lines (14), aiming to keep the endometrial model simple for tool development, our 3D *in vitro* model does not include all components of the endometrium, such as cells from the microbiome, spiral arteries, uterine glands, and the immune system (14). Additionally, due to ethical considerations, the blastocyst model employed in this study lacks an inner cell mass. Nevertheless, *ImplantoMetrics* can be adapted to include additional components (e.g., by using cells derived from primary tissues, by using more sophisticated 3D endometrial models that for instance including the immune-component of the endometrium, a different ECM composition, by inducing the window of implantation, and/or by studying receptive endometrial organoids), enabling the use of various model-specific images. Such sample-optimized 3D *in vitro* models can be directly applied in the analysis. We already demonstrated the applicability of *ImplantoMetrics* in cancer research using our model trained for BLS invasion; however, a model specifically trained for this process could provide more detailed information on the importance of involved features and their significance over time. Our model emphasizes key cell types and components essential for implantation, serving as a basis for comparison. It has to be noted that in addition to ethical restrictions, the requirement of fluorescence images of trophoblast invasion events complicates *in vivo* applications of *ImplantoMetrics* in humans.

In conclusion, *ImplantoMetrics* not only automates and enhances the accuracy of invasion analysis but also offers potential for high-throughput drug screening and personalized medicine. Future research could explore the integration of additional biological interactions to enhance the model’s predictive capabilities, especially for other 3D *in vitro* applications such as cancer experiments.

## Materials and Methods

### Cell lines and reagents

***Reagents:*** Dulbecco’s Modified Eagle’s Medium (DMEM) and McCoy’s 5A Medium were purchased from Thermo Fisher Scientific (Waltham, MA, USA), DMEM/F-12 from Invitrogen (Life Technologies, Inc., Carlsbad, CA). Heat-inactivated fetal bovine serum (FBS) was purchased from Sigma-Aldrich (St. Louis, MO, USA), the charcoal dextran-treated and heat-inactivated FBS from Gemini Bio Products (West Sacramento, CA, USA), and Matrigel from Corning (Corning, NY, USA). Additional reagents, including L-glutamine, glucose, sodium bicarbonate (3.7 g/l for DMEM-F12; Gibco), MEM non-essential amino acids, sodium pyruvate, penicillin/streptomycin, and insulin, were purchased from Fisher Scientific (Waltham, MA, USA) and Gibco (Thermo Fisher Scientific, Waltham, MA, USA) and were used in accordance with the culturing conditions for HESC, HEC-1-A, and Sw.71 cells.

***Cell lines:*** Sw.71 (21), HESC (11,16) (Accegen,Catalog No:ABI-TC257D), and HEC-1A (Elabscience Biotechnology Inc. Catalog No:EP-CL-0099) cell lines were used in the experimental studies. HESCs were cultured in DMEM medium, Sw.71 cells in DMEM-F12 medium, and HEC-1A cells in McCoy’s 5A medium with 10 µg/mL insulin supplemented. All cell types were maintained at 37°C, 95% humidity, and 5% CO_2_. Culture media were supplemented as follows: 10% fetal calf serum (FBS), 1000 U/mL penicillin, 100 µg/mL streptomycin, 10 mM HEPES, 100 nM non-essential amino acids, and 1 mM sodium pyruvate.

### Formation of blastocyst-like spheroids

BLSs were obtained following the previously described method (11). To summarize, Sw.71 cells were trypsinized and 4,000 cells were added to each well of a Costar ultra-low attachment 96-well microplate (Corning Incorporated, Corning, NY, USA). The cells were incubated for 48 hours until they formed compact spheroids, which were monitored using the Cytation 7 Plate Reader (Agilent BioTek CYT7UMW-SN). More detailed information about the cellular characteristics of trophoblast cells as a 3D model or monolayer can be found elsewhere (11).

### 3D trophoblast invasion assay

HESCs (30,000 cells) cultured in 96 well plates for 24h are covered with a Matrigel layer simulating the extracellular matrix, mixed 1:1 with DMEM (50 µl per well) After 45 minutes of waiting for the Matrigel to solidify, HEC-1-A cells (30,000 cells) are transferred on top of each well to reconstruct the epithelial component of the endometrium. This is followed by the seeding of HEC-1-A to reconstruct the epithelial component of the endometrium. A ratio of 1:1 between HESC and HEC-1-A cells was established to create ideal conditions for BLS adhesion and growth. On day four, GFP-labeled BLSs are placed on top of the cell layer. BLS invasion was monitored for up to 8 days, with images taken every 8 hours using a Cytation 7 Plate Reader (Agilent BioTek).

### 3D invasion assay with indomethacin

For the combined model, the HESC cells were prepared as described before, and then Matrigel and HEC-1-A cells were added. On the third day, three different concentrations of indomethacin from the 15 mM stock solution were added, resulting in final concentrations of 0.2 µg/ml, 0.4 µg/ml, and 0.6 µg/ml. After a 24-hour incubation period, the medium was washed with PBS and then BLS was added on top. In the models based on single cell types, cells were treated for 24 hours analogous as done in the combined model: Either HESC or HEC-1-A cells were seeded and treated for 24 hours and then washed with PBS. Matrigel and BLS were then added. These separate configurations allowed us to evaluate the specific role of each cell line in the context of invasion dynamics. Images were taken every 8 hours for five days, using a Cytation 7 plate reader and a 10x objective lens, or using an inverted fluorescence microscope (Olympus IX73). Images were analyzed using *ImplantoMetrics*.

### 3D invasion assay with TNF-α

For the 3D model, HESC cells were prepared as previously described, followed by the addition of Matrigel and HEC-1-A cells to reconstruct the epithelial component of the endometrium. On day 2, HESCs were treated with Gibco™ Human TNF-α Recombinant Protein (Thermo Fisher Scientific, Waltham, MA, USA) at three different concentrations (10 ng/ml, 25 ng/ml, and 50 ng/ml) for 24 hours. To specifically assess the effect of TNF-α, the treatment medium consisted of OptiMEM (Gibco, Thermo Fisher Scientific) mixed 1:1 with DMEM/F12 containing 1% FBS. The use of low serum concentration (1% FBS) ensured that observed effects were primarily attributable to TNF-α rather than serum-derived factors. Control wells received the same 1% FBS-containing medium without TNF-α. Following 24 hours of treatment, the medium was removed, and cells were washed three times with PBS to eliminate residual TNF-α. On day 3, Matrigel was added, and HEC-1-A cells were seeded on top to complete the epithelial reconstruction. On day 4, GFP-labeled BLSs were placed on top of the cell layers. The final medium, DMEM/F12 supplemented with 10% FBS, was used to support BLS implantation. Images were acquired every 8 hours for up to 8 days using a Cytation 7 plate reader (Agilent BioTek) with a 10x objective lens, and image analysis was performed using *ImplantoMetrics*.

### Computational tools and techniques used or analysis and visualization

Various libraries facilitated different aspects of data handling, model training, and optimization. The os library managed file paths and directories, essential for file manipulation. pandas (1.2.4) handled data manipulation and analysis, processing CSV files with image and parameter data (56,57). numpy (1.20.3) supported scientific computing with multi-dimensional arrays and mathematical functions, crucial for numerical computations (56). UMAP plots were created using the umap-learn library (version 0.5.1) to visualize high-dimensional data in a lower-dimensional space, providing insights into data clustering and structure (58). Visualizations of training and validation metrics, as well as the effects of indomethacin treatment, were created using matplotlib (3.3.4) in Python and ggplot2 in RStudio, while the datetime library managed timestamps for logging and TensorBoard callbacks (59,60). tensorflow (2.15.0) and keras (2.4.0) built, trained, and evaluated CNNs (61–63). SciPy (1.11.4) and Scikit-learn (1.2.2) performed statistical analysis and machine learning model evaluation (64,65). Hyperopt (0.2.5) optimized neural network hyperparameters (65,66). SHAP was used for model interpretability to explain predictions (67). Google.colab provided a cloud-based environment with Tesla T4 and A100 GPU availability, and its drive module enabled direct access and saving of files from Google Drive, essential for data storage and retrieval. For developing, Java 1.8, IntelliJ IDEA, and Amazon Corretto 1.8 were used. Additionally, for model conversion and layer modification, tensorflow, tf2onnx, and onnx libraries (1.10.0) were employed (35).

### Image pre-processing

***Segmentation module:*** For the segmentation module, a number of algorithms were combined to ensure separation of the target structures from the background. The rolling ball algorithm was used to mitigate unwanted background noise and uneven illumination. This algorithm interprets the image as a three-dimensional terrain, where the pixel intensities symbolize elevation. Envisioning a ball of a predefined radius “rolling” over this landscape, its contact points facilitate the estimation and subtraction of the image background (68). The effectiveness of the rolling ball algorithm in microscopy depends on selection of the radius (69). In particular, 500 µm radius was selected taking into account the morphology of the spheroids, which typically have a diameter of 200-400 µm (11). TrueColor Contrast Enhancement improves contrast without altering the original colors and shades, which is particularly important for the analysis of stained cells (70). The method proves its strength in its adaptability, responding dynamically to a variety of images and their individual characteristics. Combined with the rolling ball method for noise reduction, TrueColor Contrast Enhancement enables visibility and differentiation of cellular structures and features (70). The plugin operates in the HSB (Hue, Saturation, Brightness) color space to achieve precise and effective contrast enhancement without compromising the color integrity of the image. It avoids the challenges and limitations associated with RGB-to-HSB conversion by using a highly accurate calculation method (70). Unsharp Masking further enhances image quality by highlighting edges and details of GFP-tagged spheroids, making cell structures clearer and more distinct (71). This sequential approach uses Unsharp Masking to emphasize details while reducing excess noise and smoothing edges, further refining image quality. The result is an image with a clear and precise representation of cellular structures and features that is optimal for scientific analysis and interpretation (71).

Gaussian blurring complements the previous steps by additionally smoothing the image texture, which increases the visual quality and improves the appearance. BLS is characterized by areas of varying intensities and contrasts due to the inherent cellular and morphological complexity of the structures (11). By integrating local contrast normalization, the heterogeneous intensity distribution in GFP-labeled BLS cells is targeted, resulting in image qualities that enable precise analysis and interpretation (71). The result is superior quality images that allow precise analysis and interpretation of cellular structures and their functions, taking into account the specific intensity and contrast conditions of each region (72).

***Quality assessment of the segmentation module:*** To minimize bias and ensure the robustness of the segmentation module and its ability to differentiate between ‘Good’ and ‘Bad’ segmentations, a comprehensive quality assessment approach was implemented. ROC analysis was chosen as the primary method for evaluating image quality metrics due to its well-established application in classification problems, particularly in bioinformatics and biomedical imaging (73). The assessment was conducted using three independent datasets for each category (consisting of 13 images each, covering all time points), each segmented with *Segmentation module* images categorized as either ‘Good’ or ‘Bad’ to validate the reliability of the segmentation output. ROC curves provide a comprehensive, visual, and numerical assessment of a classifier’s ability to distinguish between ‘Good’ and ‘Bad’ images, serving as a robust framework for the pre-selection of relevant image quality metrics. This method has been extensively utilized for evaluating classification algorithms in biological sequence analysis, structural comparison, and functional annotation due to its ability to account for class imbalance and varying decision thresholds (73,74). Following ROC-based pre-selection, optimal threshold values were determined using a combination of statistical and distribution-based approaches. For metrics with a normal distribution, thresholds were set at Mean ±SD of the ‘Good’ group. For metrics displaying bimodal distributions, thresholds were defined using histogram-based analysis by selecting the midpoint between the peaks of the ‘Good’ and ‘Bad’ distributions. This combined strategy ensures that segmentation quality is assessed based on both statistical rigor and the natural separability of image quality metrics, allowing for an objective and reproducible evaluation of segmentation performance (75).

***Binary Module*:** We use the Phansalkar local thresholding method to convert grayscale images into a binary format. In our experiment, white pixels mark structures of interest—such as blastocysts and their outgrowths—while black pixels indicate the background. This process, known as binarization, simplifies the image, making it easier to measure cell features such as size, shape, and growth patterns (74). Additionally, binarization reduces image complexity, which is essential for AI model training, helping the model focus on biologically relevant patterns (76). We selected the Phansalkar method because it is designed for biological images with uneven illumination or variable fluorescence intensities—common challenges in intravital imaging (77). Unlike global thresholding methods that apply a single brightness cutoff to the entire image, Phansalkar uses local thresholds, analyzing small regions independently (77). This is particularly valuable for our data, as blastocyst outgrowths can appear faint in some areas and bright in others. The local adjustment ensures that both weak and strong signals are accurately detected, preserving crucial biological details (74,77). In our case, the window size in the Phansalkar method is the area around each pixel used to measure brightness, adjusted to match the typical size of blastocyst outgrowths. A small window captures fine details, which is important for detecting early outgrowth patterns, while a larger window smooths the image but may miss these structures. For our biological images, selecting a window size that balances detail detection and noise reduction enhances feature identification, reduces background interference, and improves reproducibility for object counting and morphological analysis (71).

### Data for training, testing, and validation of the CNN model

In developing the CNN model for automated parameter determination, we utilized a dataset comprising 1872 TIFF images of BLS invasion, captured at 0 hours and then every 8 hours up to 144 hours, ensuring an equal number of images at each interval (n=104 images per time point) using Cytation 7. These images came from successful invasion experiments without any drug treatments. The images were segmented and binarized using *ImplantoMetrics* and divided into three subsets: 80% for training, 10% for testing, and 10% for validation, while maintaining equal numbers of images per time point within the subsets. We employed transfer learning by using a pre-trained Xception model as the base (30). The final layers of this model were modified and additional layers were added to adapt it to our specific task of identifying and quantifying cell invasion metrics *in vitro*. The base layers of the Xception model remained unchanged during training, ensuring that the pre-learned features from ImageNet were retained (30,78). Data augmentation techniques such as image rotation, shearing, and mirroring were employed to increase data variance and minimize overfitting (79). The images were pre-processed using *ImplantoMetrics* to segment the input images and create binary masks. These pre-processed binary images served as input for the model, ensuring compatibility with the Xception model architecture. Min-max normalization was applied to standardize the measured data, and outliers were removed using the interquartile range (IQR) method. To further enhance the model’s performance, we employed hyperparameter optimization using the Hyperopt library (80).

***Comparison of CNN models and hyperparameter optimization:*** This process involved exploring a wide range of hyperparameters, including learning rate, number of epochs, number of layers, number of units per layer, activation functions, regularization parameters, and dropout rates *(**S7 Fig**)*. By optimizing these hyperparameters, we aimed to find the best configuration for our specific task, thus improving the model’s accuracy and robustness.

In addition to the Xception model, we compared the performance of our CNN model with other popular architectures, including ResNet, VGG16, and Inception, using ‘Hyperopt’ *(**S7 Fig**)*. Each of these architectures has unique strengths: ResNet is known for its residual connections that help in training very deep networks by mitigating the vanishing gradient problem (81); VGG16 is recognized for its simplicity and effectiveness in image classification tasks, using a straightforward architecture with small convolutional filters (81,82); Inception is known for its inception modules that allow for multi-scale feature extraction within the network (83). By evaluating these different architectures, we ensured that the chosen model, Xception (which outperformed the other models), provided the best balance of performance and computational efficiency for our task of automated parameter determination in BLS invasion images.

More specifically, the performance of these architectures was assessed based on their ability to predict the six parameters. For this purpose, the metrics MSE and MAE were used for the evaluation. MSE and MAE were chosen for the evaluation of model performance, as these are particularly meaningful in the case of normally distributed errors (84).

Minimizing the MSE and MAE is logical in such cases, as this corresponds to the models that are most likely to have generated the data, which enables an effective estimation of model accuracy (84). The learning curves for MSE and MAE offer a depiction of model fidelity and learning efficiency across 60 epochs (***Fig 4C-E***). They illustrate the model’s convergence with respect to both training and validation datasets, revealing a consistency in model performance.

***S1 Table*** presents a comparative overview of the performance of different architectures considering the training and validation metrics, which makes it possible to directly compare their effectiveness and inform model selection. Additionally, the training procedures and hyperparameters for each model are also detailed, providing a comprehensive understanding of their optimization processes **(*S7 Fig)***. ***S2 Table*** provides a detailed list of the hyperparameters used to train the Xception model, highlighting the data processing settings. ***S4 Table*** shows the stability and performance metrics of the Xception model, highlighting the robustness of the training process and the effectiveness of the model as the epochs progress.

The Xception model was ultimately selected for its superior ability to capture intricate features from the image data, leveraging depthwise separable convolutions to reduce computational cost while maintaining high accuracy (30). This makes it particularly suitable for processing the high-dimensional and complex image data used in our study.

### High dimensional feature extraction and SHAP analysis for invasion prediction

*ImplantoMetrics* employs a CNN to extract 256 high-dimensional features from images, which serve as detailed descriptors of various aspects and patterns within the data. These features are essential for accurately predicting six key parameters of the invasion process: spheroid radius, total spheroid and cell projections area, migration/invasion radius, distribution of migration/invasion, number of cell projections and circularity. The extracted features are then fed into an XGBoost algorithm, which utilizes these detailed descriptors to predict the timing of invasion events and to determine the temporal significance of each parameter. Within the CNN architecture, these features are processed through various dense layers that handle the high-dimensional data before making final predictions. To interpret the model’s predictions and understand the contribution of individual features, SHAP values were employed. SHAP analysis elucidates the dynamic significance of features over different time intervals, providing a nuanced understanding of the temporal dynamics involved in the invasion process. This integrative approach, combining feature extraction, predictive modeling, and interpretability tools, advances the study and prediction of the complex mechanisms underlying embryo implantation. By using this methodology, researchers gain a deeper understanding of which parameters are most crucial at various stages of trophoblast invasion and how their significance evolves over time.

### Image preparation with FIJI

For image analysis and model training, images were processed with FIJI, where a total of 6 parameters –spheroid radius, total spheroid and cell projections area, migration/invasion radius, distribution of migration/invasion and circularity (***S2-6 Figs***) – were analyzed separately for 1872 images. BLS radius is determined as the average dimension of height and width to reflect growth behavior and morphological diversity. The procedure for measuring the migration radius, which reflects the movement of the trophoblast cells, deals with the calculation of the average migration length and the variability of these migrations. The determination of circularity provides information about the deviation of the shape from an ideal circle. The total area and the number of trophoblast projections were quantified using the Analyze Particles tool in FIJI, providing insights into the growth dynamics and development of the BLS.

### Comprehensive evaluation and hyperparameter optimization of machine learning models for parameter importance analyze over time

We conducted a comparison of Random Forest, XGBoost, Gradient Boosting, and Support Vector Machine models, using Randomized Search for hyperparameter optimization across all models (***S3 Table***). The best performance was achieved by XGBoost based on MSE and MAE values. Using Randomized Search, parameters such as ‘colsample_bytree’, ‘learning_rate’, ‘max_depth’, ‘n_estimators’, and ‘subsample’ were tuned to enhance model performance. Validation metrics like MSE and MAE reflect the accuracy of the models. The weight of each model component in the combined model provides insights into the relative importance of each model. As ‘time’ is the main outcome that represents the process of BLS invasion in the XGBoost model, we use SHAP values to show the roles that different six parameters play over time. Each SHAP value quantifies how the inclusion or exclusion of a specific from the six parameters affect the model’s prediction, drawing upon cooperative game theory to equitably distribute the influence of each feature (67). In the construction of the model architecture, TensorFlow-Keras was employed as the foundational machine learning framework due to its comprehensive libraries and robust capabilities in training deep learning models (61). This approach highlights the specific time points at which significant shifts in the model’s predictions occur due to the collective influence of the parameters. To determine the significance of this change, we use Welch’s t-test (*p*-value ≤ 0.05), a method that can also be used with different variances.

### Analysis of lysosomal dynamics in tumor spheroids

We applied *ImplantoMetrics* to analyze lysosomal dynamics in HeLa cell tumor spheroids during apoptosis, using images from a previously published study (41). Spheroids were stained with the fluorescent probe L-lyso2 which targets lysosomes, and imaged using an Olympus laser-scanning microscope with a Mai Tai eHP Spectra Physics femtosecond laser. One-photon fluorescence images of the 3D intact tumor spheroid were captured after incubation with L-lyso, with two-photon Z-stack images taken every 2 μm section from the top to the bottom of the tumor spheroid, visualized in an 11-second video (Movie S5). From the one-photon fluorescence images of the 3D HeLa tumor spheroid incubated with L-lyso (60 µM for 1 h), we analyzed 9 images (2 µm, 10 µm, 18 µm, 26 µm, 34 µm, 42 µm, 50 µm, 58 µm, and 70 and µm) captured from a 3D Z-stack every 2 µm section from top to bottom using a 40x objective (λex = 405 nm; λem = 415–470 nm). These images were segmented and binarized using *ImplantoMetrics* to quantify the number of lysosomes, their circular distribution within the spheroid, and the total lysosomal area *(**S14 Fig)***.

### Statistics and reproducibility

All statistical analyses are described in the Methods section, with significance defined as p <0.05. Experiments included at least three biological replicates, and results are reported as mean ±SD. For the Indomethacin experiment, one-sided Welch’s t-tests were used for group comparisons, while Wilcoxon rank-sum tests were applied for the TNF-α experiment. SHAP analysis results were statistically assessed using Welch’s t-tests to determine significant differences over time. ROC curve analysis and histogram-based methods, performed using RStudio (version 4.2.2), were employed for threshold determination during segmentation. All other statistical analyses were conducted using Python (version 3.11) with SciPy (version 1.11.4).

## Data availability

All images used to assess the efficacy of indomethacin on BLS were uploaded to GitHub (https://github.com/creativebrain1729/ImplantoMetrics/releases/tag/v.1.0.0) as ‘Negative.control-Indomethacin-.model.zip’. Additional images of control experiments were uploaded as ‘exampleimages-control.zip’, and images for segmentation quality assessment as ‘Quality_assesment_datsets-1.zip’. Underlying data for images were uploaded as excel files.

## Code availability

The Fiji plugin *ImplantoMetrics* developed for our study is available on GitHub (https://github.com/creativebrain1729/ImplantoMetrics/releases/tag/v.1.0.0). Installation instructions and documentation for the plugin are provided in the respective GitHub repositories.

## Acknowledgements

We would like to express our deepest gratitude to Nejat Güneş, who provided valuable assistance in preparing the images for CNN training. His encouragement was very valuable throughout the research process.

## Author contributions

**Conceptualization**: Ayberk Alp Gyunesh, Marlene Rezk-Füreder, Barbara Arbeithuber.

**Methodology:** Ayberk Alp Gyunesh, Marlene Rezk-Füreder, Patrick Stelzl, Barbara Arbeithuber.

**Software:** Ayberk Alp Gyunesh.

**Investigation:** Ayberk Alp Gyunesh

**Data curation:** Ayberk Alp Gyunesh, Celine Kapper.

**Formal analysis:** Ayberk Alp Gyunesh.

**Visualization**: Ayberk Alp Gyunesh.

**Project administration:** Marlene Rezk-Füreder, Barbara Arbeithuber.

**Supervision:** Marlene Rezk-Füreder, Barbara Arbeithuber.

**Validation:** Ayberk Alp Gyunesh, Celine Kapper.

**Resources:** Gil Mor.

**Writing – original draft:** Ayberk Alp Gyunesh, Marlene Rezk-Füreder, Barbara Arbeithuber.

**Writing – review & editing:** Ayberk Alp Gyunesh, Marlene Rezk-Füreder, Celine Kapper, Gil Mor, Omar Shebl, Peter Oppelt, Patrick Stelzl, Barbara Arbeithuber.

## Supporting information overview

### Supplementary Figures

**S1 Fig. Comparison of CNN architectures and hyperparameter optimization with ‘’hyperopt’’.**

**S2 Fig. Calculation of the spheroid radius.**

**S3 Fig. Calculation of the migration/invasion radius.**

**S4 Fig. Calculation of distribution of migration/invasion.**

**S5 Fig. Calculation of circularity.**

**S6 Fig. Calculation of total spheroid and cell projections area and number of cell projections.**

**S7 Fig. Dynamic SHAP value analysis of significant CNN features over time intervals.**

**S8 Fig. Workflow for installing and using ImplantoMetrics in Fiji/ImageJ.**

**S9 Fig. Quality assessment of image segmentation.**

**10 Fig. Evaluation of segmentation quality using ROC analysis.**

**S11 Fig. Evaluation of segmentation quality using histogram-based thresholding.**

**S12 Fig. Impact of indomethacin on the invasion factor measured with the Olympus fluorescence microscope.**

**S13 Fig. Impact of TNF-α on the Invasion factor.**

**S14 Fig. Analysis of lysosomal dynamics in HeLa cells using *ImplantoMetrics*.**

### Supplementary Tables

**S1 Table. Evaluation of model accuracy and effectiveness.**

**S2 Table. Hyperparameters for Xception model training.**

**S3 Table. Performance metrics and hyperparameters for parameter/feature importance analysis.**

**S4 Table. Xception model training and performance metrics.**

