## Supplementary Information for "*ImplantoMetrics* - Multidimensional trophoblast invasion assessment by combining 3D-*in-vitro* modeling and deep learning analysis"

**Supplementary Figures** **2**

S1 Figure. Comparison of CNN architectures and hyperparameter optimization with ‘’hyperopt’’. 2
S2 Figure. Calculation of the spheroid radius. 3
S3 Figure. Calculation of the migration/invasion radius. 4
S4 Figure. Calculation of distribution of migration/invasion. 5
S5 Figure. Calculation of circularity. 6
S6 Figure. Calculation of total spheroid and cell projections area and number of cell projections. 7
S7 Figure. Dynamic SHAP value analysis of significant CNN features over time intervals. 8
S8 Figure. Workflow for installing and using ImplantoMetrics in Fiji/ImageJ. 9
S10 Figure. Evaluation of segmentation quality using ROC analysis. 11
S11 Figure . Evaluation of segmentation quality using histogram-based thresholding. 12
S12 Figure. Impact of indomethacin on the invasion factor measured with the Olympus fluorescence microscope. 13
Supplementary Figure 13. Impact of TNF-α on the Invasion factor. 14
S14 Figure. Analysis of lysosomal dynamics in HeLa cells using ImplantoMetrics 15

**Supplementary Tables** **16**

S1 Table. Evaluation of model accuracy and effectiveness. 16
S2 Table. Hyperparameters for Xception model training. 16
S3 Table. Performance metrics and hyperparameters for parameter/feature importance analysis. 18
S4 Table. Xception model training and performance metrics. 19

**References** **20**

### Supplementary Figures

**S1 Figure. Comparison of CNN architectures and hyperparameter optimization with ‘‘hyperopt”.**

**(A)** Performance of the VGG16 model, including training and validation metrics for Mean Squared Error (MSE) and Mean Absolute Error (MAE). The hyperparameters used for this model are listed, indicating the configuration that was tested. Despite its widespread use, VGG16 exhibited higher error rates, suggesting it was less suited for our specific application. **(B)** Performance of the Resnet50 model, displaying the training and validation MSE and MAE metrics. The corresponding hyperparameters for this model are detailed, showing the parameters used during optimization. Resnet50 demonstrated improved performance over VGG16, but there were still noticeable fluctuations in the error rates, indicating potential for further optimization. **(C)** Performance of the InceptionV3 model. **(D)** Hyperparameter search space utilized for optimizing all models. The search space included a variety of parameters such as learning rate, number of epochs, number of layers, units per layer, activation functions, and regularization techniques. **(E)** Performance of the Xception model, which included training and validation metrics for MSE , MAE and Loss. The Xception model, based on HyperOpt analysis, demonstrated the best performance among all the models evaluated. It showed consistently lower error rates and more stable performance, making it the most suitable model for our specific application.

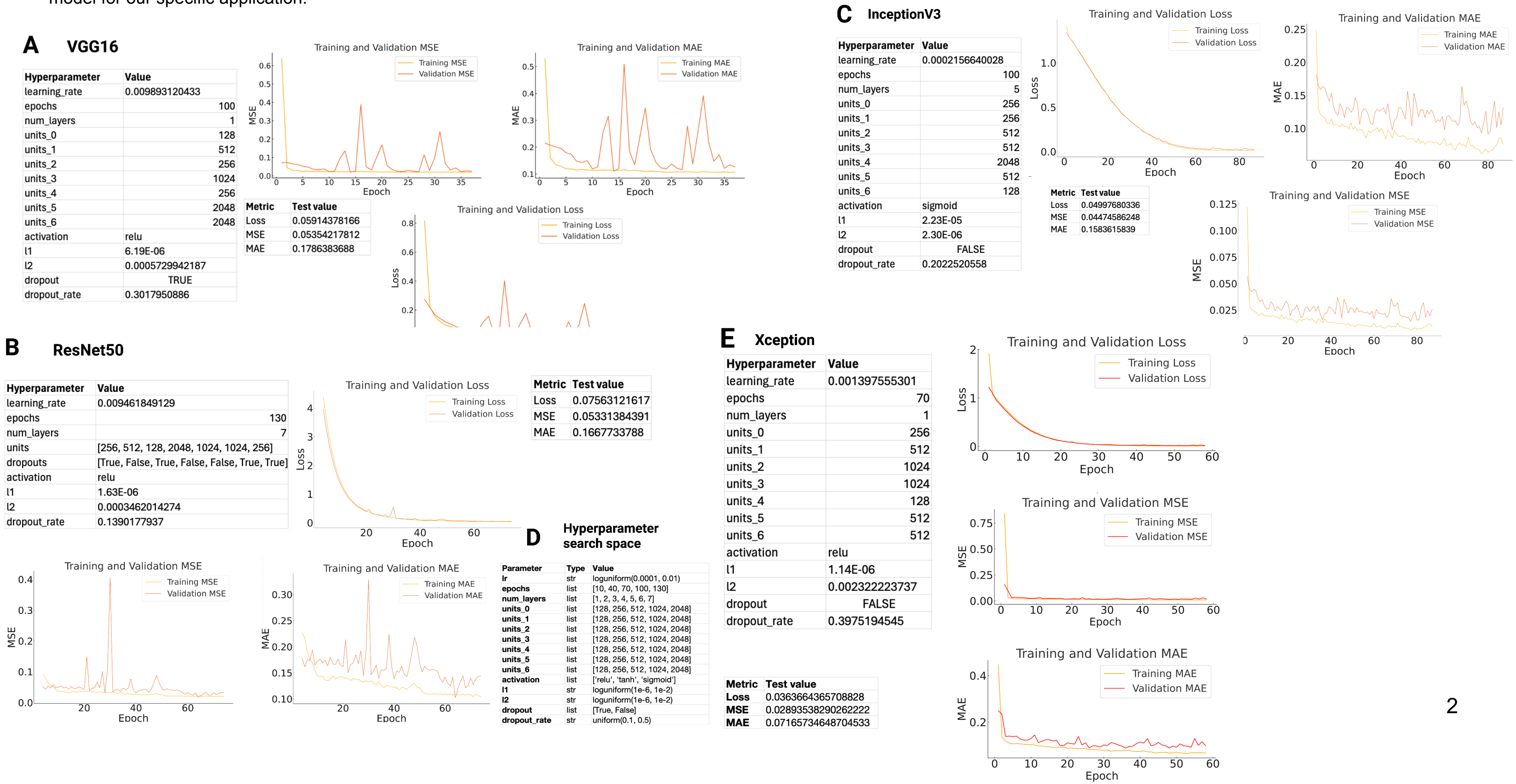

**A VGG16**

| Hyperparameter | Value |
|---|---|
| learning_rate | 0.009893120433 |
| epochs | 100 |
| num_layers | 1 |
| units_0 | 128 |
| units_1 | 512 |
| units_2 | 256 |
| units_3 | 1024 |
| units_4 | 256 |
| units_5 | 2048 |
| units_6 | 2048 |
| activation | relu |
| l1 | 6.19E-06 |
| l2 | 0.0005729942187 |
| dropout | TRUE |
| dropout_rate | 0.3017950886 |

| Metric | Test value |
|---|---|
| Loss | 0.05914378166 |
| MSE | 0.05354217812 |
| MAE | 0.1786383688 |

**B ResNet50**

| Hyperparameter | Value |
|---|---|
| learning_rate | 0.009461849129 |
| epochs | 130 |
| num_layers | 7 |
| units | [256, 512, 128, 2048, 1024, 1024, 256] |
| dropouts | [True, False, True, False, False, True, True] |
| activation | relu |
| l1 | 1.63E-06 |
| l2 | 0.0003462014274 |
| dropout_rate | 0.1390177937 |

| Metric | Test value |
|---|---|
| Loss | 0.07563121617 |
| MSE | 0.05331384391 |
| MAE | 0.1667733788 |

**C InceptionV3**

| Hyperparameter | Value |
|---|---|
| learning_rate | 0.0002156640028 |
| epochs | 100 |
| num_layers | 5 |
| units_0 | 256 |
| units_1 | 256 |
| units_2 | 512 |
| units_3 | 512 |
| units_4 | 2048 |
| units_5 | 512 |
| units_6 | 128 |
| activation | sigmoid |
| l1 | 2.23E-05 |
| l2 | 2.30E-06 |
| dropout | FALSE |
| dropout_rate | 0.2022520558 |

| Metric | Test value |
|---|---|
| Loss | 0.04997680336 |
| MSE | 0.04474586248 |
| MAE | 0.1583615839 |

**D Hyperparameter search space**

| Parameter | Type | Value |
|---|---|---|
| **lr** | str | loguniform(0.0001, 0.01) |
| **epochs** | list | [10, 40, 70, 100, 130] |
| **num_layers** | list | [1, 2, 3, 4, 5, 6, 7] |
| **units_0** | list | [128, 256, 512, 1024, 2048] |
| **units_1** | list | [128, 256, 512, 1024, 2048] |
| **units_2** | list | [128, 256, 512, 1024, 2048] |
| **units_3** | list | [128, 256, 512, 1024, 2048] |
| **units_4** | list | [128, 256, 512, 1024, 2048] |
| **units_5** | list | [128, 256, 512, 1024, 2048] |
| **units_6** | list | [128, 256, 512, 1024, 2048] |
| **activation** | list | ['relu', 'tanh', 'sigmoid'] |
| **l1** | str | loguniform(1e-6, 1e-2) |
| **l2** | str | loguniform(1e-6, 1e-2) |
| **dropout** | list | [True, False] |
| **dropout_rate** | str | uniform(0.1, 0.5) |

**E Xception**

| Hyperparameter | Value |
|---|---|
| learning_rate | 0.001397555301 |
| epochs | 70 |
| num_layers | 1 |
| units_0 | 256 |
| units_1 | 512 |
| units_2 | 1024 |
| units_3 | 1024 |
| units_4 | 128 |
| units_5 | 512 |
| units_6 | 512 |
| activation | relu |
| l1 | 1.14E-06 |
| l2 | 0.002322223737 |
| dropout | FALSE |
| dropout_rate | 0.3975194545 |

| Metric | Test value |
|---|---|
| **Loss** | 0.0363664365708828 |
| **MSE** | 0.02893538290262222 |
| **MAE** | 0.07165734648704533 |

**S2 Figure. Calculation of the spheroid radius.**
To ensure a reliable and representative measurement of the average size of a spheroid (such as a BLS) without projections, the radius (R) was determined with the oval selection tool by averaging the height (H) and width (W). This method reflects an average dimension that takes into account both the growth behavior and the morphological diversity of the spheroid in the culture dish. Because spheroids can vary in shape, the average provides a standardized measure that compensates for individual deviations from perfect spherical shape, providing a robust basis for quantitative comparisons and further statistical analysis.

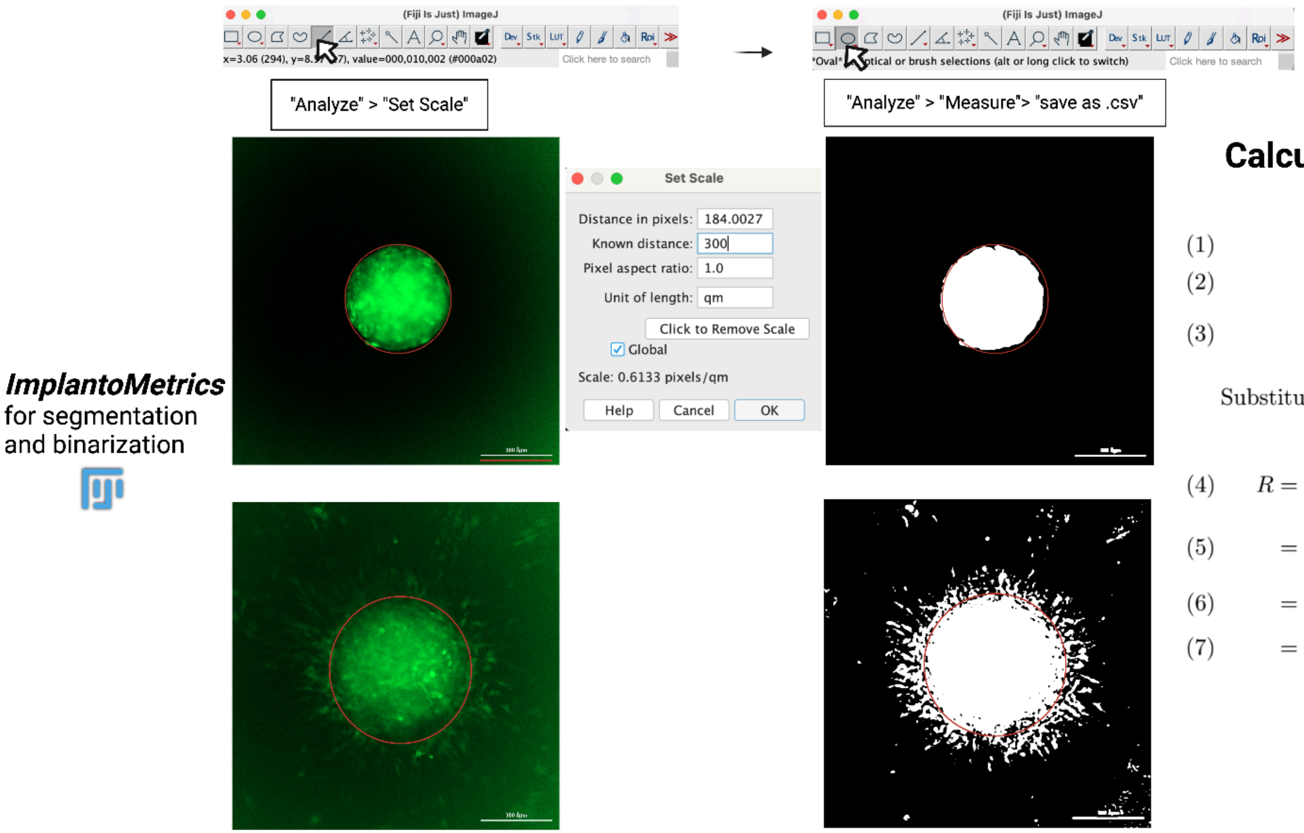

| | Label | Width | Height |
|---|---|---|---|
| 1 | sample.tif | 4,337E+02 | 4,451E+02 |

### Calculation of spheroid radius

(1) $$W = 4.337 \times 10^2 \quad \text{(Width in units)}$$

(2) $$H = 4.451 \times 10^2 \quad \text{(Height in units)}$$

(3) $$R = \frac{1}{2}\left(\frac{W}{2} + \frac{H}{2}\right) \quad \text{(Average radius)}$$

Substituting the values for W and H, we obtain:

(4) $$R = \frac{1}{2}\left(\frac{4.337 \times 10^2}{2} + \frac{4.451 \times 10^2}{2}\right)$$

(5) $$= \frac{1}{2}(216.85 + 222.55)$$

(6) $$= \frac{1}{2}(439.4)$$

(7) $$= 219.7$$

**S3 Figure. Calculation of the migration/invasion radius.**
In the analysis of cell migration, the use of vector summation provides a measure of displacement from a central origin point. Each vector represents the trajectory of an individual cell or particle, with the magnitude corresponding to the distance migrated. For untreated cells, which serve as a control group, the assumption is that the movements are random and independent. By summing these vectors, we compile the aggregate movement of the cell population. The magnitude of the resultant vector quantifies the overall migratory activity away from the origin.

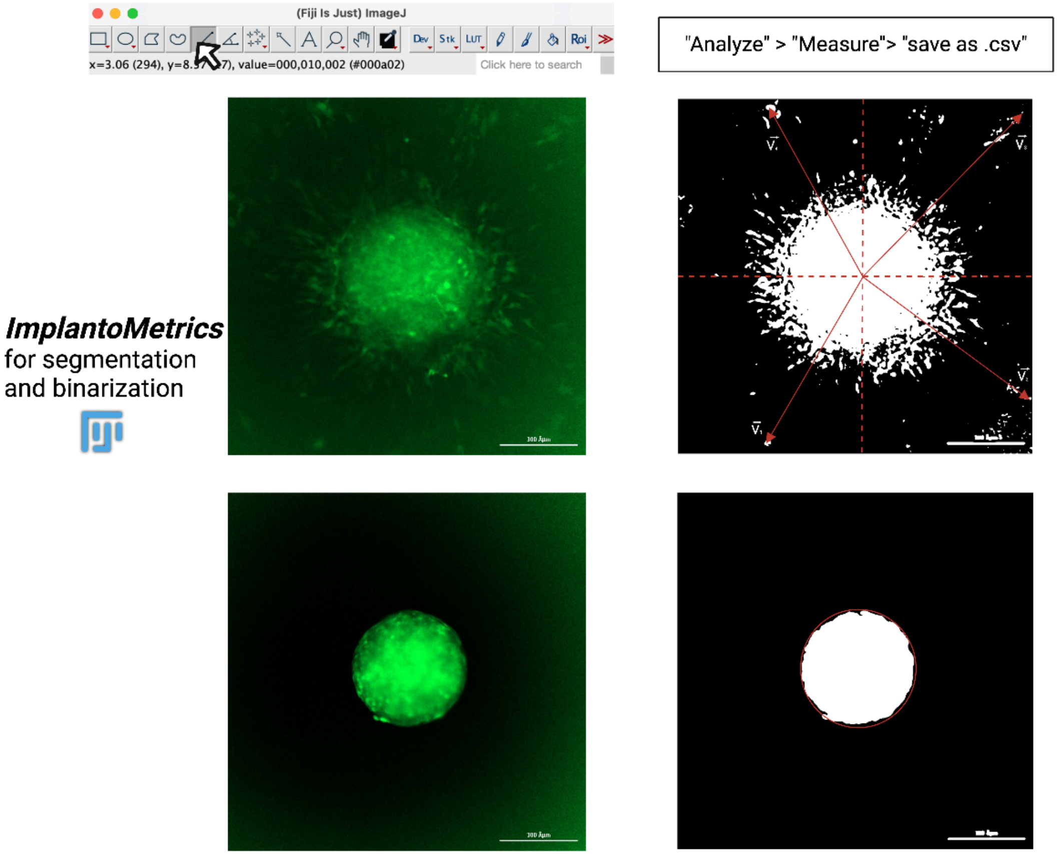

| | Label | X | Y | Length |
|---|---|---|---|---|
| 1 | vector1.tif | 5,118E+02 | 3,545E+02 | 7,252E+02 |
| 2 | vektor2.tif | 1,017E+03 | 4,348E+02 | 7,799E+02 |
| 3 | vektor3.tif | 9,995E+02 | 9,762E+02 | 8,816E+02 |
| 4 | vektor4.tif | 5,214E+02 | 9,842E+02 | 7,577E+02 |

→ the long migration radius

→ Coordinates

### Calculation of migration/invasion radius

$$\vec{V_1} = 511.8\hat{\imath} + 354.5\hat{\jmath} \qquad \text{Angle: } 34.71^\circ$$
$$\vec{V_2} = 1017.0\hat{\imath} + 434.8\hat{\jmath} \qquad \text{Angle: } 23.15^\circ$$
$$\vec{V_3} = 999.5\hat{\imath} + 976.2\hat{\jmath} \qquad \text{Angle: } 44.32^\circ$$
$$\vec{V_4} = 521.4\hat{\imath} + 984.2\hat{\jmath} \qquad \text{Angle: } 62.09^\circ$$

The summation of the vectors $(\vec{V_1} + \vec{V_2} + \vec{V_3} + \vec{V_4})$ yields:

$$\text{Sum Magnitude (Length)} = 4106.28$$
$$\text{Sum X-Component} = 3049.7$$
$$\text{Sum Y-Component} = 2749.7$$
$$\text{Summed Angle (in Degrees)} = 42.04^\circ$$

The vector notation for the sum is:

$$\vec{V}_{\text{Sum}} = 3049.7\hat{\imath} + 2749.7\hat{\jmath}$$

**S4 Figure. Calculation of distribution of migration/invasion.**
The mean length is an average measure of the extent of cell migration, while the variance and standard deviation provide information about the variability of these migrations. A higher standard deviation indicates a greater dispersion of migration distances, suggesting heterogeneity within the migration behavior of the cell. Conversely, a lower standard deviation would indicate a more uniform migration pattern.

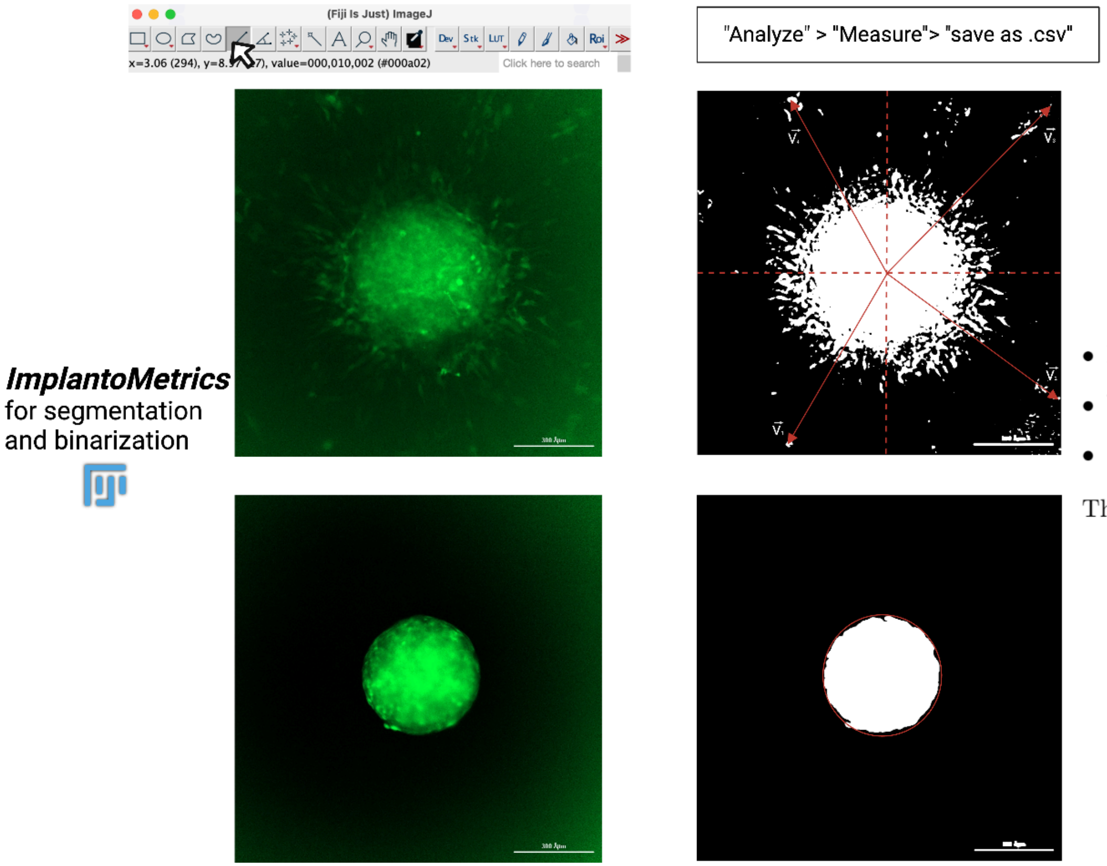

| | Label | X | Y | Length |
|---|---|---|---|---|
| 1 | vector1.tif | 5,118E+02 | 3,545E+02 | 7,252E+02 |
| 2 | vektor2.tif | 1,017E+03 | 4,348E+02 | 7,799E+02 |
| 3 | vektor3.tif | 9,995E+02 | 9,762E+02 | 8,816E+02 |
| 4 | vektor4.tif | 5,214E+02 | 9,842E+02 | 7,577E+02 |

→ the long migration radius

→ Coordinates

### Calculation of distribution of migration/invasion

- Mean length of migration/invasion radius ($\mu$): 786.1
- Variance of migration/invasion radius lengths ($\sigma^2$): 3418.51
- Standard Deviation of migration/invasion radius lengths ($\sigma$): 58.47

The calculation of the variance is performed using the formula:

$$\sigma^2 = \frac{1}{n}\sum_{i=1}^{n}(L_i - \mu)^2$$

**S5 Figure. Calculation of circularity.**
A circularity value of 1 indicates a perfect circle. As the shape deviates from a circular form, the circularity value becomes less than 1, suggesting an increasingly elongated or irregular shape. The formula emphasizes the relationship between the area of the object and the square of its perimeter, with the constant 4 pi scaling the value such that a circle yields a circularity of 1 [1].

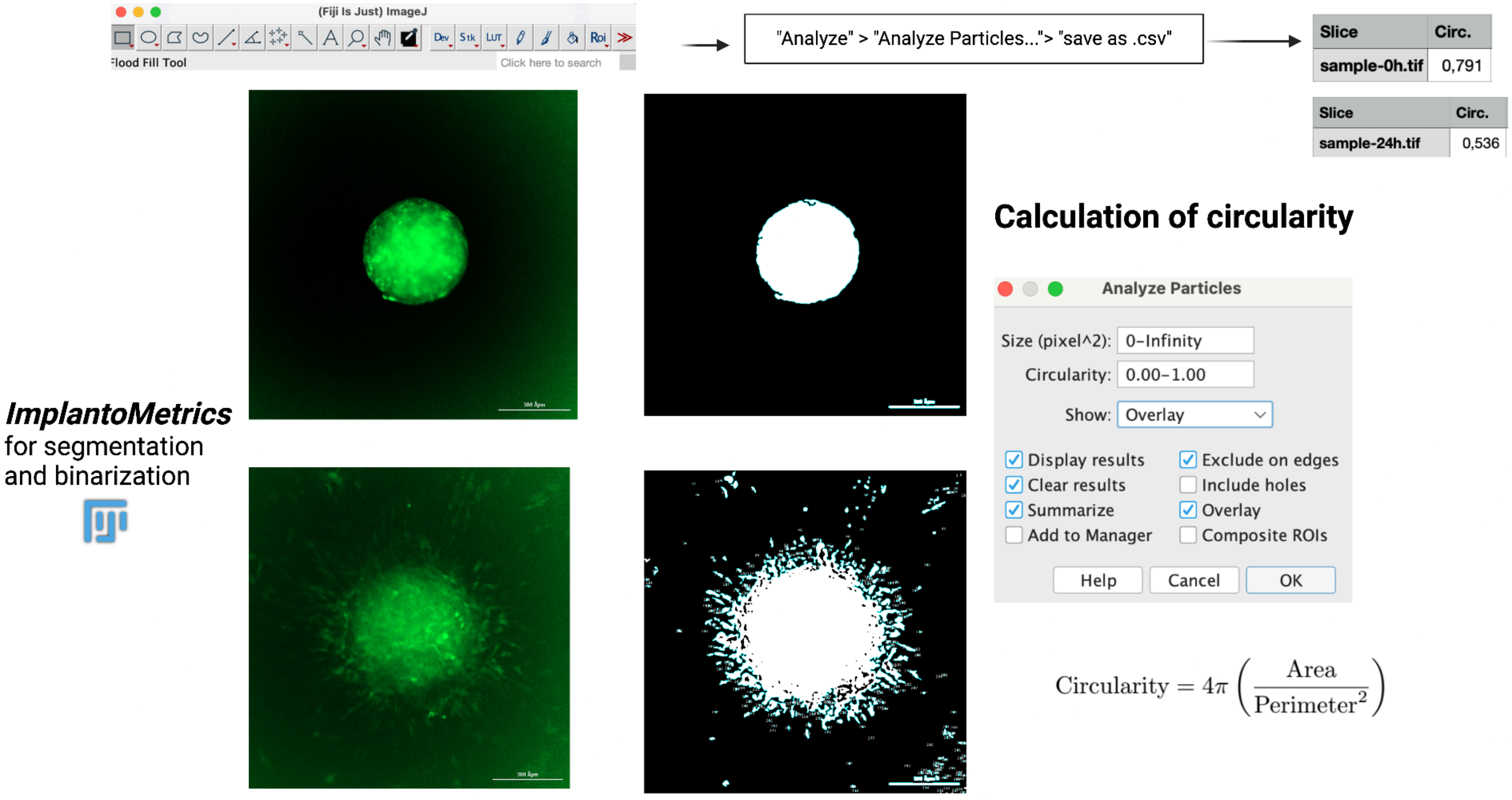

**S6 Figure. Calculation of total spheroid and cell projections area and number of cell projections.**
The figure demonstrates the method for quantifying the total area and number of trophoblast projections in blastocyst-like spheroids (BLS) at two time points: 0 and 24 hours. The left side displays fluorescence images of BLS, prepared for segmentation and binarization using Implantometrics. Following binarization, as shown in the middle, the BLS and its projections are distinctly visible. The right side presents a table with the quantitative analysis results: 'sample-0h.tif' shows a single main cell with no projections, whereas 'sample-24h.tif' shows a significant increase in total area and number of projections, indicating growth and development of projections. The total area is measured in square micrometers (μm²), and the number of projections is determined by subtracting 1 from the total particle count, excluding the main cell as the largest particle.

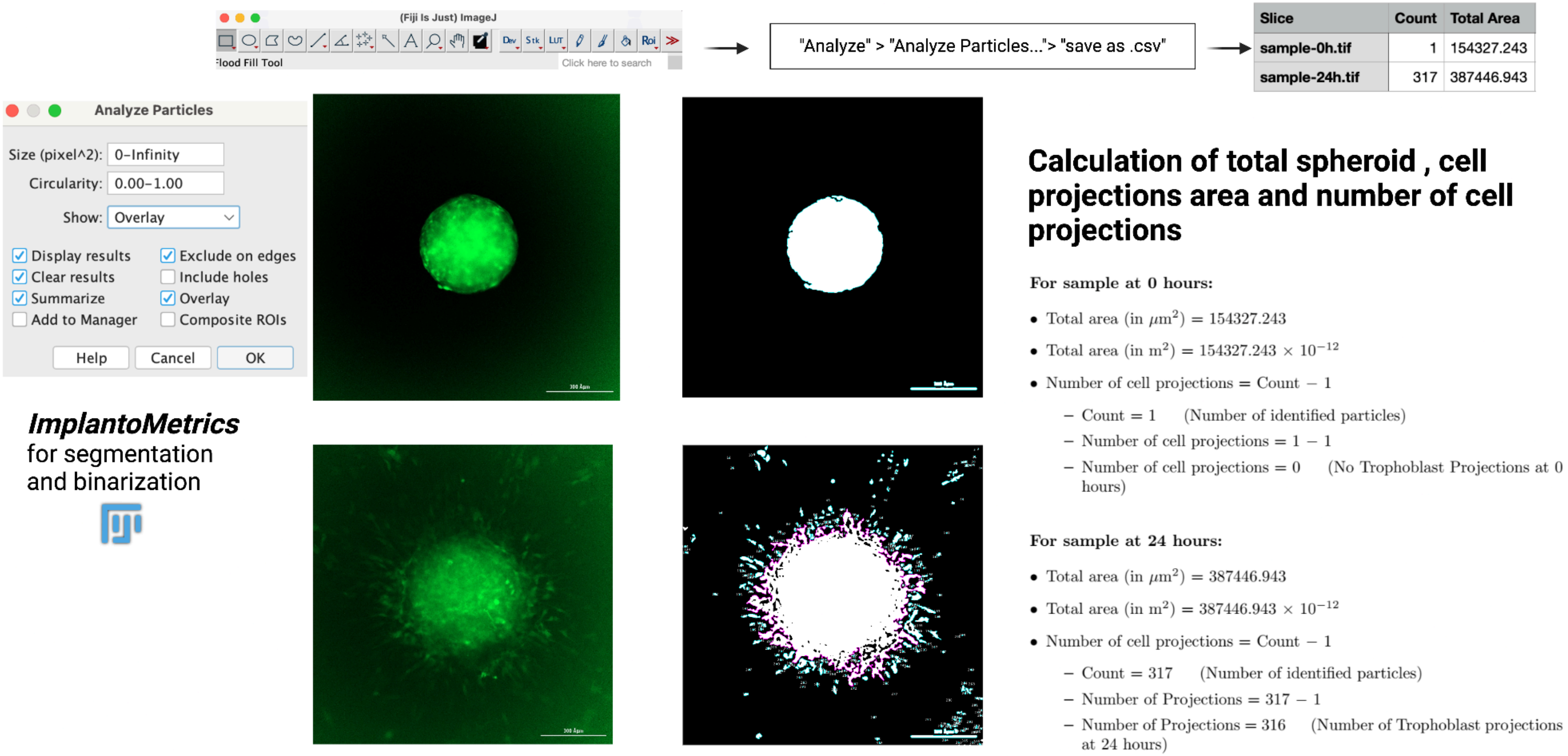

**S7 Figure. Dynamic SHAP value analysis of significant CNN features over time intervals.** Detailed analysis of the SHAP values, highlighting the importance of each significant CNN feature over different time intervals. The plots show the SHAP values for five different time intervals: 32-40 hours, 48-56 hours, 64-80 hours, 80-96 hours and 96-144 hours post-BLS-placement. The characteristics are listed on the y-axis, with those with the greatest effect at the top. The x-axis represents the SHAP value, which indicates the change in model output ‘’time’’ when that feature is present. Each point in the plot represents the SHAP value of a feature for an observation. The color of the points indicates the actual values of the feature: blue for low values and red for high values. Looking at the plots, it is clear that some features are consistently important across time intervals, while others vary in importance. This variation highlights the dynamic nature of the invasion process and the need to develop models that account for this temporal variability. The p-values reported for each time window confirm the statistical significance of the observed differences in feature importance. A low p-value (≤ 0.05) indicates that the observed differences in SHAP scores across time intervals are statistically significant, supporting the hypothesis that feature importance changes over time.

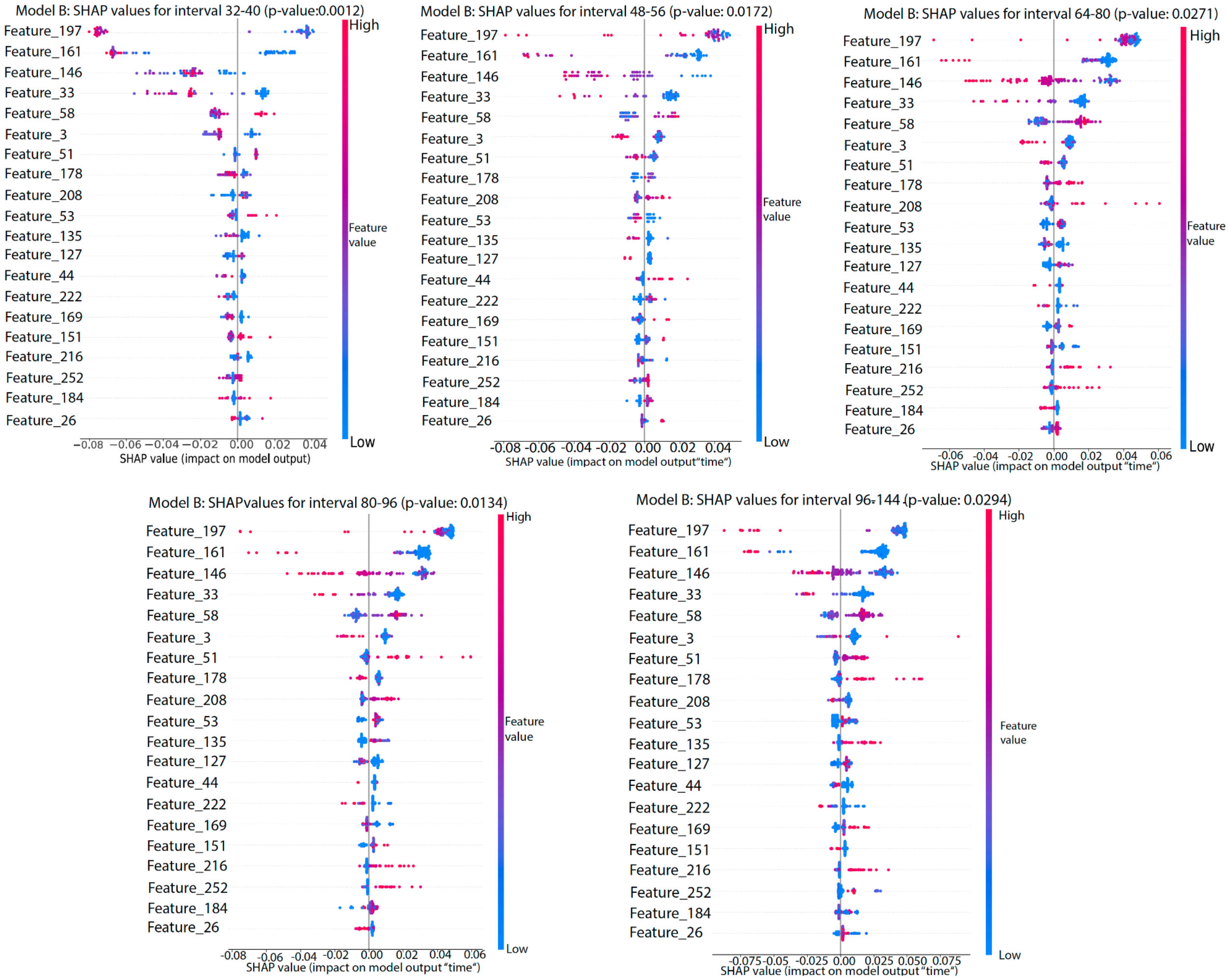

**S8 Figure. Workflow for installing and using *ImplantoMetrics* in Fiji/ImageJ.**

This figure illustrates the step-by-step workflow for installing and using *ImplantoMetrics* within Fiji/ImageJ for fluorescence image analysis. (1) First, Fiji/ImageJ must be downloaded from its official website (https://imagej.net/software/fiji/downloads), ensuring compatibility with the user's operating system (Windows, macOS, or Linux). (2) The Fiji plugin *ImplantoMetrics* is available on GitHub (https://github.com/creativebrain1729/ImplantoMetrics/releases/tag/v.1.0.0). After installation, the *ImplantoMetrics* plugin is added by locating the Fiji installation folder, copying the `.jar` plugin file into the plugins subdirectory, and restarting Fiji to activate the module. (3) Additionally, the BioVoxxel Toolbox is required for preprocessing functions and can be installed either through Fiji's built-in updater or via manual download from its repository. (4) Once the installation is complete, fluorescence images can be imported into Fiji in TIFF or PNG format, ensuring that the object of interest (e.g., spheroids or blastocyst-like structures) is clearly visible in a single fluorescence channel, typically GFP. (5) Users can proceed with *ImplantoMetrics* by following a structured workflow for segmentation and analysis (see Figure 2 for details).

### 1 Download Fiji/ImageJ

1-Go to **https://imagej.net/software/fiji/**

2-**Download:** Choose the version appropriate for your operating system (Windows, macOS, Linux).

Downloads

| ~ Download Fiji for your OS ~ | |
|---|---|
| Windows 64-bit | imagej.net (USA), micron.ox.ac.uk (European mirror) |
| Windows 32-bit | imagej.net (USA), micron.ox.ac.uk (European mirror) |
| macOS (x86_64) | imagej.net (USA), micron.ox.ac.uk (European mirror) |
| Linux (64-bit) | imagej.net (USA), micron.ox.ac.uk (European mirror) |
| No JRE | imagej.net (USA), micron.ox.ac.uk (European mirror) |

### 2 Install the *ImplantoMetrics* Plugin

**1-Locate your FIJI installation folder:**
- After unzipping or installing FIJI, you typically have a folder named Fiji.app.

**2-Find the plugins folder:**
- Inside the Fiji.app folder, there is a subfolder called "plugins".

**3-Copy the plugin file (ImplantoMetrics_.jar):**
- Download or obtain the ImplantoMetrics_.jar file
- https://github.com/creativebrain1729/ImplantoMetrics/releases/tag/v.1.0.0
- Move (or copy) this file into the Fiji.app/plugins directory.

**4-Restart FIJI:**
- If FIJI was open, close it completely.
- Then reopen Fiji to load the newly added plugin.

### 3 Install the BioVoxxel Toolbox

**Option A: Using Fiji's built-in updater**

Launch FIJI.
Go to Help → Update....
Click Manage Update Sites.
Scroll down to find and check the box next to BioVoxxel.
Click Close, then Apply Changes.
Restart FIJI when prompted.

**Option B: Manual download**

Visit the BioVoxxel Toolbox GitHub page (e.g. https://github.com/BioVoxxel/BioVoxxel-Toolbox/releases).
Download the latest .jar release.
Copy the .jar file into Fiji.app/plugins.
Restart FIJI.

### 4 Open Your Fluorescence Images

1-**Launch Fiji** by double-clicking the ImageJ-win64.exe (or your system's equivalent) inside the Fiji.app folder.
Import your images:

2-Go to the top menu and select **File → Open**.

3-Choose the fluorescence images you wish to analyze (e.g., TIFF or PNG files).

4-The images should be in a format where the object of interest (e.g., spheroid, blastocyst-like structure 10x or 20x objective) is clearly visible in one channel (often green for GFP).

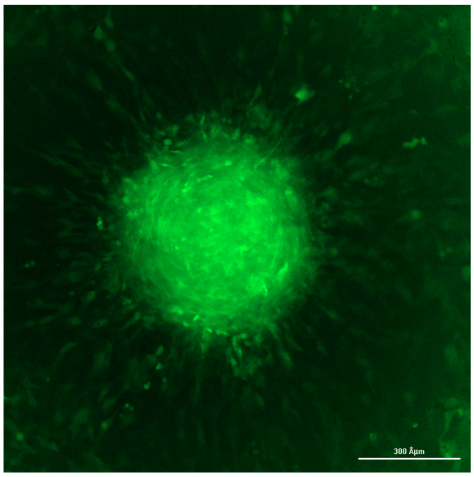

### 5 Run *ImplantoMetrics* (Figure 2)

**S9 Figure. Quality assessment of image segmentation.**

The Segmentation Module is used to preprocess fluorescence images. This step includes contrast enhancement, unsharp masking, and Gaussian blurring to optimize image quality. The rolling ball algorithm is applied to remove background noise, ensuring a clearer visualization of the spheroid structure. Once the segmentation process is complete, users will receive feedback on whether the segmentation was successful or unsuccessful based on a quality assessment algorithm. If segmentation is successful, users can proceed directly to the Binary Module, where Phansalkar's local thresholding method is applied to further refine the segmented image. If segmentation is unsuccessful, troubleshooting suggestions based on image parameters such as entropy, contrast, local entropy variance, median intensity, and SNR are provided, guiding users on how to adjust settings in Fiji to improve segmentation quality.

### Successful Segmentation ✅

**Segmentation was successful**

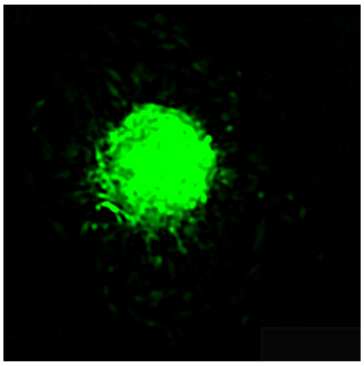

- Clear and well-defined boundaries.
- Accurate identification of the spheroid and cell projections.
- Minimal background noise or artifacts.
-
- The segmented image corresponds well with the original fluorescence image.

**No further adjustments needed. The segmentation is ready for binary module.**

### Unsuccessful Segmentation & Troubleshooting ❌

**Segmentation was not successful troubleshooting required**

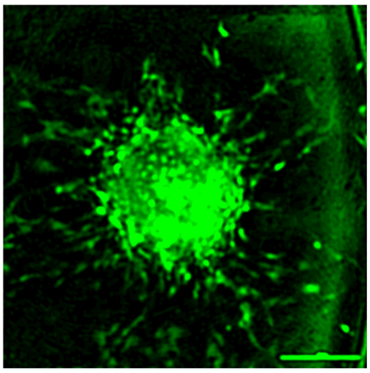

**Troubleshooting Table for FIJI Based on High-AUC (>0.80) Parameters and Thresholds e.g.**

| Parameter | Issue Detected | Threshold | Suggested Fix in FIJI |
|---|---|---|---|
| Mean Intensity (too low) | Image too dark, poor signal detection | 25.90 | Increase exposure or use: *Process > Enhance Contrast...* |
| SNR (too low) | High noise, weak signal clarity | 1.57 | Use: *Process > Noise > Despeckle or Gaussian Blur...* |
| Laplacian Variance (too high) | Excessive texture, high noise | 432.56 | Apply smoothing: *Process > Filters > Gaussian Blur...* |
| Skewness (too high) | Image asymmetry, inconsistent intensity patterns | 3.47 | Adjust brightness uniformity: *Process > Subtract Background...* |

**S10 Figure. Evaluation of segmentation quality using ROC analysis.**
Three independent datasets of segmented images, each containing both "Good" and "Bad" segmentations, were used for this analysis to ensure robustness and generalizability. ROC curves for selected image quality metrics are shown, illustrating their classification performance in distinguishing between "Good" and "Bad" images. Metrics with an AUC greater than 0.80 were selected for further analysis, including Mean, Standard Deviation (SD), Signal-to-Noise Ratio (SNR), Laplacian Variance, Histogram Spread, Median Intensity, Skewness, Kurtosis, Variance of Local Entropy, Angular Second Moment, Contrast, Correlation, Entropy, Homogeneity, and Contrast Range.

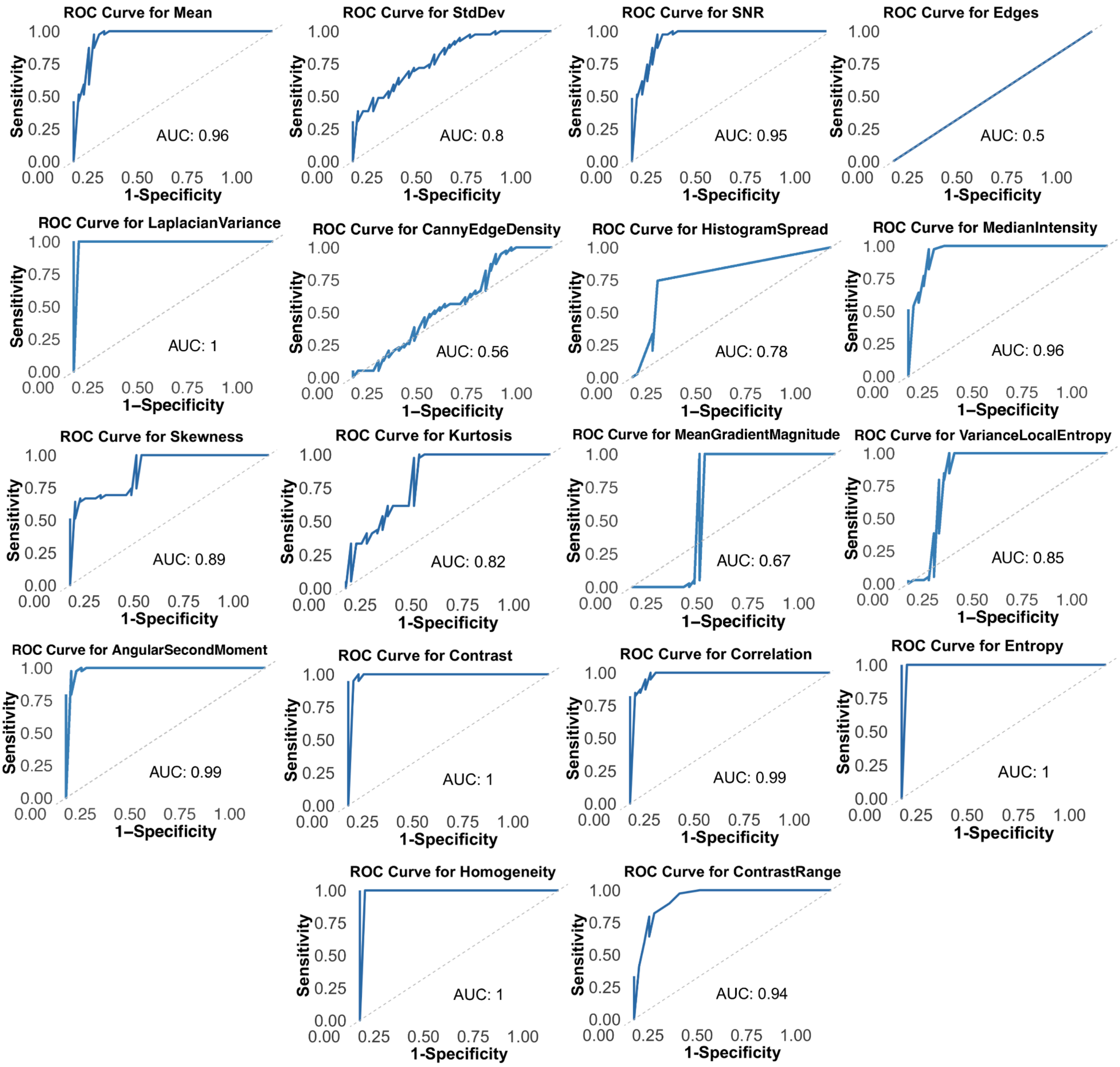

**S11 Figure . Evaluation of segmentation quality using histogram-based thresholding.**
Three independent datasets of segmented images, each containing both "Good" and "Bad" segmentations, were used for this analysis to ensure robustness and generalizability. Histograms of the selected metrics are shown, with the threshold values indicated by a dashed blue line. The thresholds were determined using a combination of two statistical approaches: first, the Mean ± Standard Deviation (SD) of the "Good" group was used to account for natural variations in image quality, assuming a normal distribution; second, for metrics displaying bimodal distributions, the threshold was set at the midpoint between the peaks of the "Good" and "Bad" groups to better capture the natural separation in data.

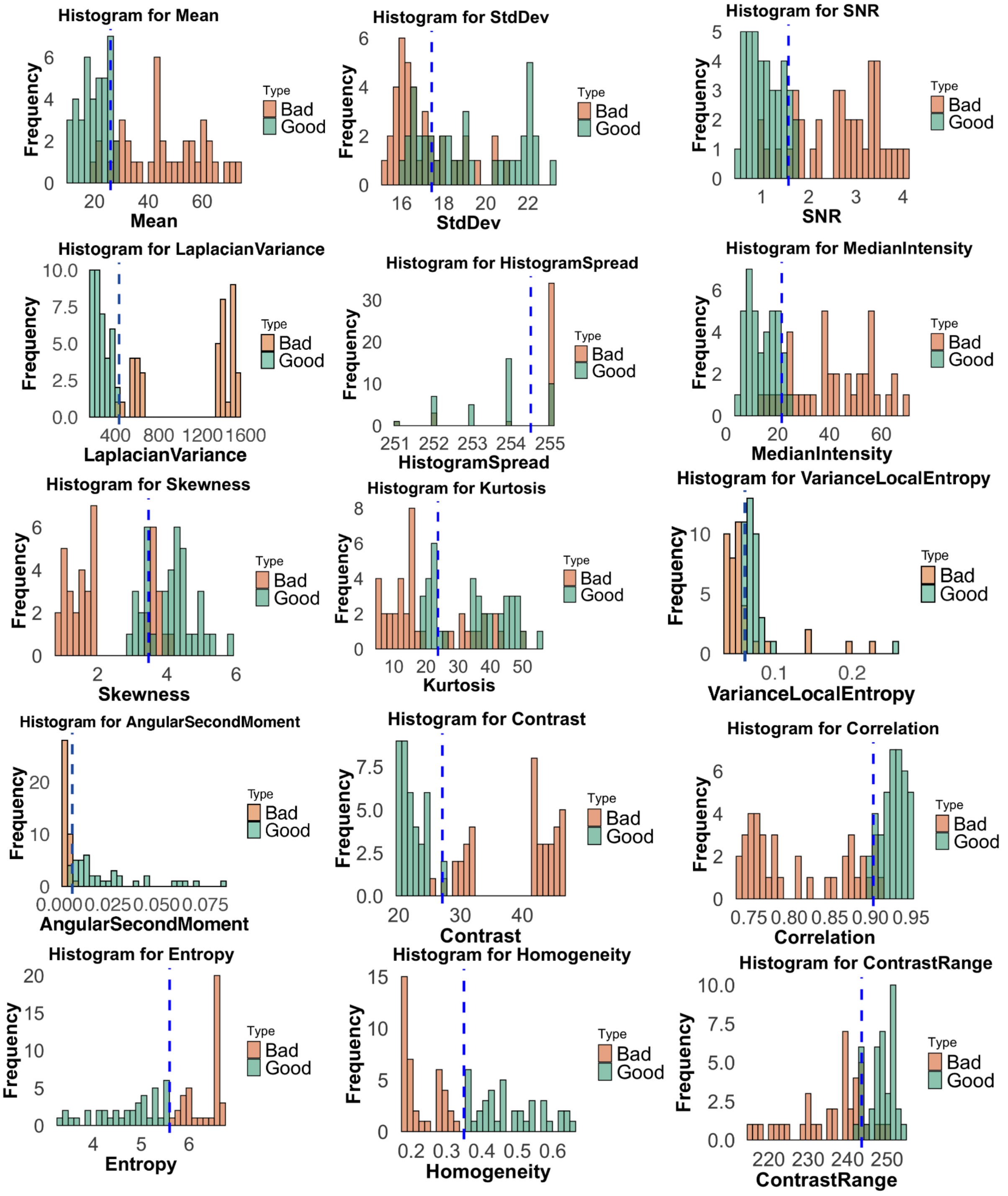

**S12 Figure. Impact of indomethacin on the invasion factor measured with the Olympus fluorescence microscope.**
**(A)** Combined model system, (**B)** only HESC with Matrigel, and (**C)** only HEC-1-A with Matrigel. The blue curves represent the untreated cells, the orange curves show the treatment with 0.2 µg/ml, the green line with 0.4 µg/ml, and the red line with 0.6 µg/ml indomethacin. In all figures, the presence of a single asterisk symbolizes statistically significant differences with p-values ≤ 0.05, as determined by Welch's t-test.

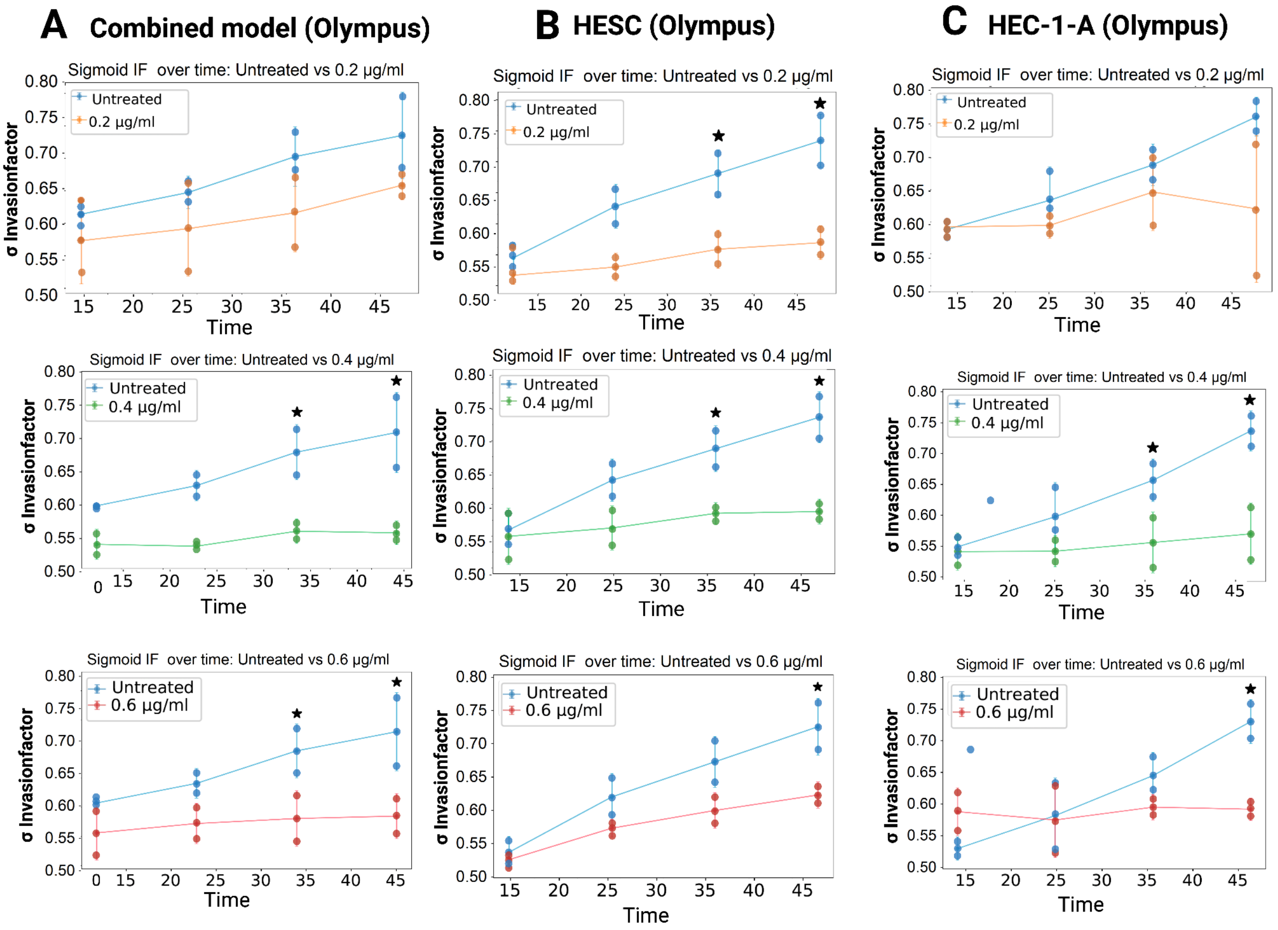

**S13 Figure. Impact of TNF-α on the Invasion factor.**

Time-dependent changes in the *Invasion factor* were assessed for three experimental conditions (n=3, distinct samples per treatment condition) using combined model (HESC, Matrigel, and HEC-1-A): **(A)** untreated vs. 10 ng/ml TNF-α, **(B)** untreated vs. 25 ng/ml TNF-α, and **(C)** untreated vs. 50 ng/ml TNF-α. The blue curve represents the treatment with 10 ng/ml TNF-α, the green curve indicates treatment with 25 ng/ml TNF-α, and the red curve shows treatment with 50 ng/ml TNF-α. Statistically significant differences ($p \leq 0.05$) are indicated by asterisks, as determined by Wilcoxon rank-sum test (two-sided). Error bars represent the standard deviation (± SD).

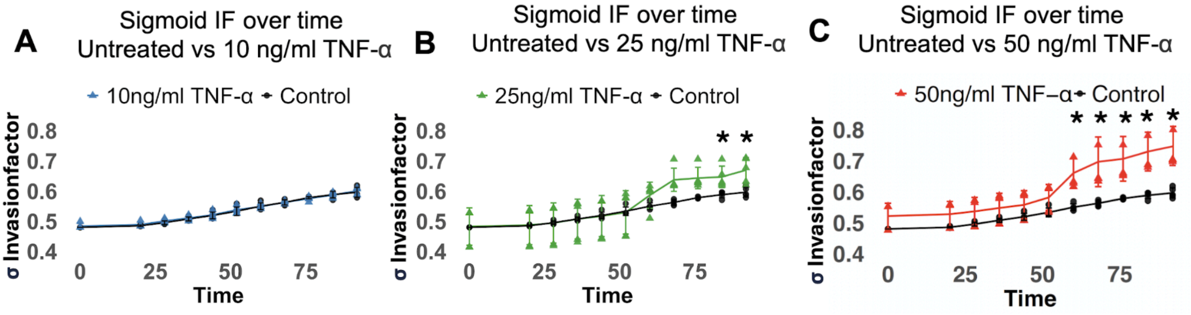

**S14 Figure. Analysis of lysosomal dynamics in HeLa cells using *ImplantoMetrics.***
This figure demonstrates the versatility of *ImplantoMetrics* in analyzing lysosomal dynamics in HeLa cell tumor spheroids stained with the fluorescent probe L-lyso [2]. **(A)** I*mplantoMetrics* processed the images (n=9, measured at 2, 10, 18, 26, 34, 42, 50, 58, and 70 µm), **(B)** binarized them, and analyzed parameters such as circularity of the lysosomal distribution in the spheroid, number of lysosomes, and total area of lysosomes. **(C)** The analysis quantified lysosomal number, circularity, and total area. (**D**) Circularity of the lysosomal distribution in the spheroid in dependence of the measured spheroid position. (**E**) The number of lysosomes in dependence of the measured spheroid position. **(F)** Total area of lysosomes in dependence of the measured spheroid position.

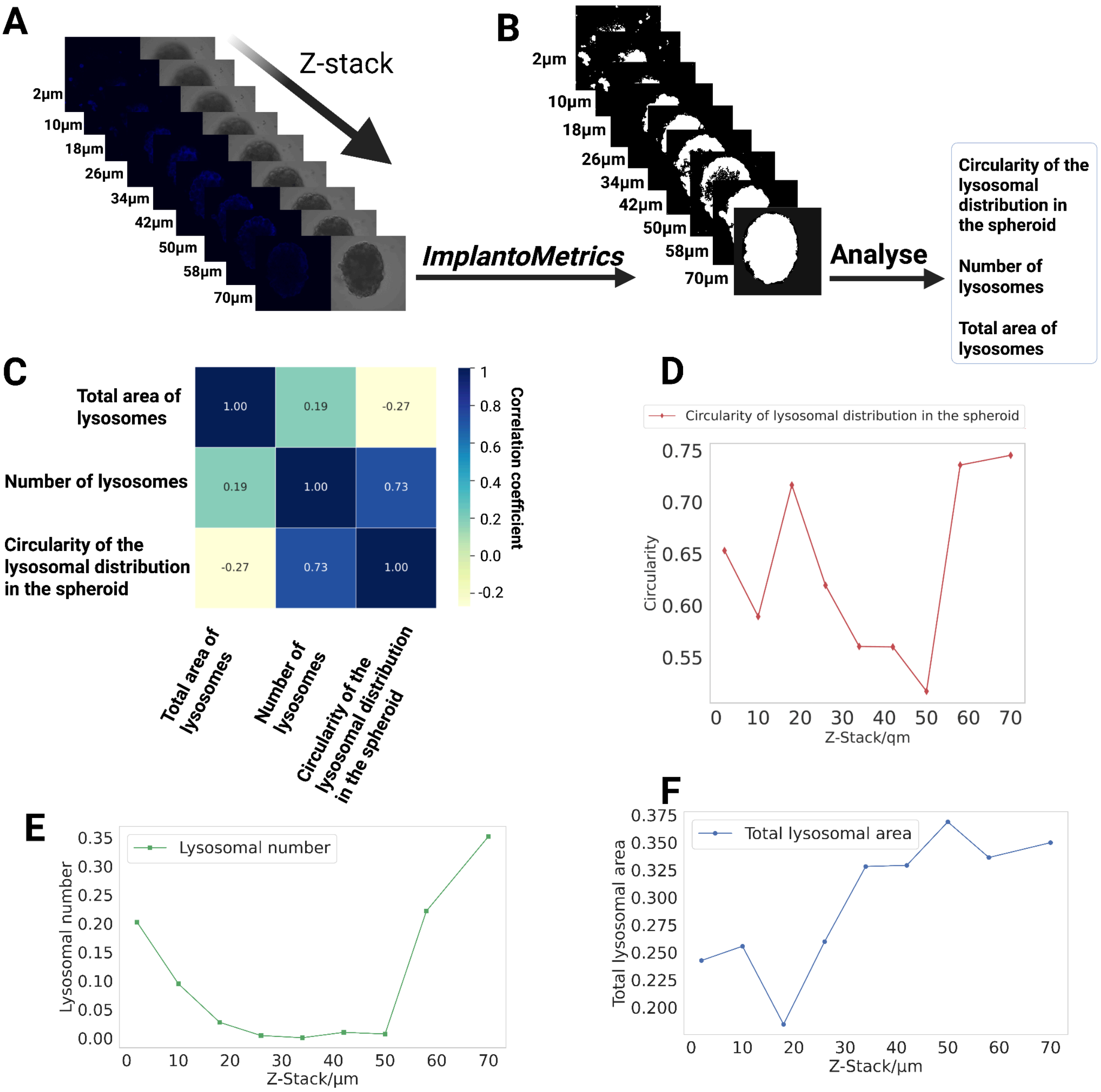

### Supplementary Tables

**S1 Table. Evaluation of model accuracy and effectiveness.**

Performance comparison of different CNN architectures, including VGG16, Xception, ResNet50 and InceptionV3. Training and validation metrics such as Loss, MSE and MAE are compared. Xception consistently shows the best performance across all measured metrics in both training and validation. In particular, the lower validation loss and error measures should be emphasized, indicating that the Xception model is more effective in this specific application than the other models tested.

| Model | Training Loss | Training MSE | Training MAE | Validation Loss | Validation MSE | Validation MAE |
|---|---|---|---|---|---|---|
| **VGG16** | 0.02430 | 0.020305 | 0.10746 | 0.02880 | 0.02487 | 0.126678 |
| **Xception** | 0.00976 | 0.008487 | 0.06666 | 0.01880 | 0.01666 | 0.068379 |
| **ResNet50** | 0.03621 | 0.01923 | 0.10351 | 0.04965 | 0.03279 | 0.14444 |
| **InceptionV3** | 0.01374 | 0.00916 | 0.20510 | 0.07574 | 0.02563 | 0.13121 |

**S2 Table. Hyperparameters for Xception model training.**

The table presents a list of hyperparameters used to train the Xception model. These include data processing settings such as rotation range and shear intensity, which increase the diversity of the training data, as well as parameters such as batch size and target size, which influence the model and the training process. In addition, the number of epochs and the initial learning rate are specified, which determine the duration and speed of the training. Other parameters such as dropout rate and kernel regularizer are used to avoid overfitting, while the choice of optimizer and activation functions influences the efficiency of the training. This detailed list of hyperparameters makes it possible to reproduce and understand the training process of the model.

| Hyperparameter | Value | Explanation |
|---|---|---|
| **learning_rate** | 0.001397 | Determines the step size during model training. Smaller values lead to slower but potentially better learning. |
| **epochs** | 70 | Number of complete passes through the training dataset. More epochs can improve accuracy but risk overfitting. |
| **num_layers** | 1 | Number of layers in the model. More layers can capture complex patterns but require more computation. |
| **units_0** | 256 | Number of neurons in layer 0. More neurons increase model capacity but also computational cost. |
| **units_1** | 512 | Number of neurons in layer 1. More neurons increase model capacity but also computational cost. |
| **units_2** | 1024 | Number of neurons in layer 2. More neurons increase model capacity but also computational cost. |

| | | |
|---|---|---|
| **units_3** | 1024 | Number of neurons in layer 3. More neurons increase model capacity but also computational cost. |
| **units_4** | 128 | Number of neurons in layer 4. More neurons increase model capacity but also computational cost. |
| **units_5** | 512 | Number of neurons in layer 5. More neurons increase model capacity but also computational cost. |
| **units_6** | 512 | Number of neurons in layer 6. More neurons increase model capacity but also computational cost. |
| **activation** | relu | Activation function introducing non-linearity, allowing the model to learn complex patterns. ReLU is commonly used. |
| **l1** | 1.14e-06 | L1 regularization to prevent overfitting by penalizing large weights, leading to a sparse model. |
| **l2** | 0.002322223737 | L2 regularization to prevent overfitting by penalizing large weights, keeping many weights small. |
| **dropout** | FALSE | Indicates if dropout is used. Dropout is a technique to prevent overfitting by randomly setting units to 0. |
| **dropout_rate** | 0.3975194545 | Fraction of units to drop if dropout is used. Not utilized as dropout is set to FALSE. |
| **rescale** | 1/255 | Scaling factor to normalize pixel values to the range [0, 1]. |
| **rotation_range** | 45 | Range for random rotations during data augmentation, in degrees. |
| **width_shift_range** | 0.2 | Range for random horizontal shifts during data augmentation, as a fraction of total width. |
| **height_shift_range** | 0.2 | Range for random vertical shifts during data augmentation, as a fraction of total height. |
| **shear_range** | 0.2 | Range for random shearing transformations during data augmentation. |
| **horizontal_flip** | TRUE | Whether to randomly flip images horizontally during data augmentation. |
| **fill_mode** | 'nearest' | Mode to fill in newly created pixels during transformations. |
| **Batch Normalization** | Used | Normalization technique to stabilize and speed up training by adjusting and scaling the activations. |

**S3 Table. Performance metrics and hyperparameters for parameter/feature importance analysis.**

| XGBoost (eXtreme Gradient Boosting) | | | |
|---|---|---|---|
| | **Model A** | **Model B** | **Combined Model** |
| **MSE** | 0.001852 | 0.002838 | 0.0056 |
| **MAE** | 0.012087 | 0.021702 | 0.0541 |
| **RMSE** | 0.015482 | 0.039095 | 0.0751 |
| **colsample_bytree** | 0.7673 | 0.7438 | |
| **gamma** | 0.0 | 0.0 | |
| **learning_rate** | 0.01 | 0.0236 | |
| **max_depth** | 8.0 | 5.0 | |
| **n_estimators** | 933.0 | 100.0 | |
| **reg_alpha** | 2.0 | 0.0 | |
| **reg_lambda** | 8.0 | 1.0 | |
| **subsample** | 0.5 | 0.8437 | |
| **Weight** | 0.394875 | 0.605125 | 0.4169 |
| **Model A Normalized Weight** | | | 0.394875 |
| **Model B Normalized Weight** | | | 0.605125 |

| Random Forest | | | |
|---|---|---|---|
| | **Model A** | **Model B** | **Combined Model** |
| **MSE** | 0.0267 | 0.0087 | 0.0107 |
| **MAE** | 0.1278 | 0.0749 | 0.0818 |
| **RMSE** | 0.1635 | 0.0934 | 0.1035 |
| **colsample_bytree** | 0.9129 | 0.6 | |
| **gamma** | 0.0 | 0.0 | |
| **learning_rate** | 0.3 | 0.01 | |
| **max_depth** | 6.0 | 3.0 | |
| **n_estimators** | 1000.0 | 1000.0 | |
| **reg_alpha** | 2.0 | 0.0 | |
| **reg_lambda** | 8.0 | 1.0 | |
| **subsample** | 0.6 | 0.6 | |
| **Weight** | 0.1524 | 0.4672 | 0.3804 |
| **Model A Normalized Weight** | | | 0.246 |
| **Model B Normalized Weight** | | | 0.754 |

| Gradient Boosting | | | |
|---|---|---|---|
| | **Model A** | **Model B** | **Combined Model** |
| **MSE** | 0.0146 | 0.0062 | 0.0072 |
| **MAE** | 0.0946 | 0.0595 | 0.0681 |
| **RMSE** | 0.1210 | 0.0791 | 0.0848 |
| **max_depth** | 10.0 | 10.0 | |
| **min_samples_leaf** | 2.0 | 1.0 | |
| **min_samples_split** | 2.0 | 2.0 | |
| **n_estimators** | 1000.0 | 461.0 | |
| **Weight** | 0.1861 | 0.4351 | 0.3787 |
| **Model A Normalized Weight** | | | 0.2996 |
| **Model B Normalized Weight** | | | 0.7003 |

| Super Vector Machine | | | |
|---|---|---|---|
| | **Model A** | **Model B** | **Combined Model** |
| **MSE** | 0.0157 | 0.0054 | 0.0056 |
| **MAE** | 0.0996 | 0.0409 | 0.0541 |
| **RMSE** | 0.1251 | 0.0737 | 0.0751 |
| **learning_rate** | 0.01 | 0.01 | |
| **max_depth** | 10 | 9 | |
| **max_features** | log2 | auto | |
| **n_estimators** | 433 | 904 | |
| **subsample** | 0.6 | 0.6 | |
| **Weight** | 0.1503 | 0.4328 | 0.4169 |
| **Model A Normalized Weight** | | | 0.2577 |
| **Model B Normalized Weight** | | | 0.7423 |

**S4 Table. Xception model training and performance metrics.**

Performance indicators of the Xception model during the training process. It includes metrics such as training and validation loss, MSE and MAE to demonstrate the stable learning process and the effectiveness of the model. The metrics show that the model is neither overfitting nor underfitting, which is confirmed by the consistent approximation of the training and validation values.

| Epoch | Training Loss | Validation Loss | Training MSE | Validation MSE | Training MAE | Validation MAE |
|---|---|---|---|---|---|---|
| 1 | 1,916 | 1,222 | 0,840 | 0,158 | 0,446 | 0,249 |
| 2 | 1,064 | 1,093 | 0,031 | 0,097 | 0,135 | 0,234 |
| 3 | 0,981 | 0,947 | 0,024 | 0,032 | 0,118 | 0,138 |
| 4 | 0,893 | 0,864 | 0,021 | 0,035 | 0,109 | 0,140 |
| 5 | 0,808 | 0,776 | 0,020 | 0,032 | 0,106 | 0,137 |
| 6 | 0,725 | 0,696 | 0,021 | 0,033 | 0,108 | 0,140 |
| 7 | 0,646 | 0,616 | 0,020 | 0,028 | 0,106 | 0,127 |
| 8 | 0,572 | 0,542 | 0,019 | 0,024 | 0,105 | 0,122 |
| 9 | 0,505 | 0,478 | 0,019 | 0,023 | 0,103 | 0,122 |
| 10 | 0,445 | 0,424 | 0,019 | 0,026 | 0,102 | 0,130 |
| 11 | 0,391 | 0,378 | 0,018 | 0,031 | 0,101 | 0,145 |
| 12 | 0,342 | 0,324 | 0,017 | 0,021 | 0,097 | 0,112 |
| 13 | 0,301 | 0,287 | 0,017 | 0,023 | 0,097 | 0,118 |
| 14 | 0,263 | 0,253 | 0,017 | 0,024 | 0,096 | 0,124 |
| 15 | 0,230 | 0,222 | 0,016 | 0,023 | 0,093 | 0,124 |
| 16 | 0,201 | 0,196 | 0,015 | 0,023 | 0,091 | 0,119 |
| 17 | 0,176 | 0,174 | 0,015 | 0,024 | 0,090 | 0,122 |
| 18 | 0,155 | 0,146 | 0,015 | 0,017 | 0,091 | 0,099 |
| 19 | 0,136 | 0,129 | 0,015 | 0,017 | 0,091 | 0,099 |
| 20 | 0,119 | 0,117 | 0,015 | 0,019 | 0,088 | 0,110 |
| 21 | 0,105 | 0,104 | 0,015 | 0,020 | 0,089 | 0,107 |
| 22 | 0,093 | 0,090 | 0,014 | 0,018 | 0,087 | 0,105 |
| 23 | 0,081 | 0,093 | 0,014 | 0,030 | 0,086 | 0,138 |
| 24 | 0,073 | 0,071 | 0,014 | 0,017 | 0,087 | 0,102 |
| 25 | 0,065 | 0,067 | 0,014 | 0,020 | 0,087 | 0,111 |
| 26 | 0,056 | 0,055 | 0,012 | 0,014 | 0,081 | 0,089 |
| 27 | 0,050 | 0,052 | 0,012 | 0,017 | 0,081 | 0,098 |
| 28 | 0,045 | 0,046 | 0,013 | 0,016 | 0,082 | 0,097 |
| 29 | 0,041 | 0,041 | 0,012 | 0,015 | 0,080 | 0,093 |
| 30 | 0,036 | 0,039 | 0,012 | 0,017 | 0,077 | 0,101 |
| 31 | 0,033 | 0,037 | 0,012 | 0,017 | 0,080 | 0,099 |
| 32 | 0,029 | 0,037 | 0,011 | 0,020 | 0,074 | 0,112 |
| 33 | 0,027 | 0,039 | 0,011 | 0,024 | 0,077 | 0,108 |
| 34 | 0,025 | 0,031 | 0,011 | 0,019 | 0,077 | 0,102 |
| 35 | 0,023 | 0,027 | 0,011 | 0,016 | 0,076 | 0,096 |
| 36 | 0,022 | 0,024 | 0,012 | 0,015 | 0,078 | 0,088 |
| 37 | 0,020 | 0,024 | 0,011 | 0,015 | 0,078 | 0,095 |
| 38 | 0,018 | 0,024 | 0,010 | 0,017 | 0,074 | 0,096 |
| 39 | 0,017 | 0,025 | 0,010 | 0,019 | 0,074 | 0,107 |
| 40 | 0,016 | 0,026 | 0,011 | 0,021 | 0,074 | 0,114 |
| 41 | 0,016 | 0,026 | 0,011 | 0,022 | 0,076 | 0,110 |
| 42 | 0,015 | 0,032 | 0,010 | 0,028 | 0,072 | 0,129 |
| 43 | 0,014 | 0,020 | 0,010 | 0,016 | 0,071 | 0,098 |
| 44 | 0,012 | 0,022 | 0,009 | 0,019 | 0,067 | 0,102 |
| 45 | 0,013 | 0,024 | 0,009 | 0,021 | 0,068 | 0,112 |
| 46 | 0,012 | 0,021 | 0,009 | 0,019 | 0,070 | 0,101 |
| 47 | 0,012 | 0,018 | 0,009 | 0,016 | 0,069 | 0,098 |
| 48 | 0,011 | 0,017 | 0,009 | 0,014 | 0,067 | 0,088 |
| 49 | 0,011 | 0,020 | 0,009 | 0,018 | 0,068 | 0,097 |
| 50 | 0,012 | 0,017 | 0,010 | 0,014 | 0,071 | 0,092 |
| 51 | 0,011 | 0,021 | 0,009 | 0,019 | 0,067 | 0,106 |
| 52 | 0,010 | 0,019 | 0,008 | 0,017 | 0,066 | 0,098 |
| 53 | 0,010 | 0,018 | 0,008 | 0,016 | 0,065 | 0,096 |
| 54 | 0,010 | 0,018 | 0,009 | 0,016 | 0,068 | 0,091 |
| 55 | 0,010 | 0,029 | 0,009 | 0,027 | 0,068 | 0,130 |
| 56 | 0,010 | 0,019 | 0,009 | 0,018 | 0,068 | 0,100 |
| 57 | 0,010 | 0,030 | 0,009 | 0,029 | 0,068 | 0,115 |
| 58 | 0,010 | 0,019 | 0,008 | 0,018 | 0,067 | 0,098 |
